# A Single-Cell Atlas Of Human Pediatric Liver Reveals Age-Related Hepatic Gene Signatures

**DOI:** 10.1101/2025.04.16.649149

**Authors:** Rachel D Edgar, Diana Nakib, Damra Camat, Sai Chung, Patricia Lumanto, Jawairia Atif, Catia T. Perciani, Xue-Zhong Ma, Cornelia Thoeni, Nilosa Selvakumaran, Justin Manuel, Blayne Sayed, Koen Huysentruyt, Amanda Ricciuto, Ian McGilvray, Yaron Avitzur, Gary D Bader, Sonya A MacParland

**Author notes:** corresponding **Address for Correspondence** (R.D. Edgar), (Y. Avitzur), (G.D. Bader), (S.A. MacParland). co-first authors (Rachel D Edgar, Diana Nakib, Damra Camat). co-senior authors (Yaron Avitzur, Gary D Bader, Sonya A MacParland). **Author contributions** Conceptualization: RDE, YA, GDB, SAM Formal analysis: RDE, DN, SC, PL, CT Investigation: RDE, DN, DC, SC, PL, JA, CTP, XM, JM Resources: BS, KH, AR, IM, YA, GDB, SAM Funding acquisition: IM, YA, GDB, SAM Project administration: NS Writing – original draft: RDE Writing – review & editing: All Authors.

## Abstract

**Background & Aims:** The liver plays a critical role in metabolism and immune function, yet the contributions of its heterogeneous cell types to these processes remain unclear. While most liver studies focus on adults, pediatric liver diseases often present differently, underscoring the need for age-specific research.

**Approach & Results:** To better understand cellular drivers of childhood liver diseases, we generated single-cell RNA-seq (scRNA-seq) maps of the normal pediatric liver and used this map to examine disease-related populations in biopsies from pediatric patients with Intestinal Failure-Associated Liver Disease (IFALD). The normal pediatric liver map consists of 42,660 cells from 9 donors aged 2-17 years. Compared to normal adult liver (26,372 cells; 7 donors, age 26-69) pediatric livers exhibited differences in myeloid populations. Specifically, pediatric Kupffer-like cells (*MARCO*+*C1QA*+*VSIG4*+) exhibited higher expression of immune activation genes, including *CCL4, CCL3* and *IL1B. In vitro* stimulation confirmed more IL1-β secreting myeloid cells in pediatric versus adult livers, supporting these findings. Using the pediatric atlas as a reference, we analyzed three IFALD biopsies (11,969 cells; 3 donors, ages 4 months–9 years) and identified increased expression of fibrosis-associated genes (e.g., *LY96*) in Kupffer-like cells. Additionally, mesenchymal cells in IFALD showed fibrotic gene modules resembling adult liver cells more than healthy pediatric cells. These signatures, undetectable when comparing IFALD to adult liver alone, highlighting the value of a pediatric map.

**Conclusions:** Taken together, our healthy pediatric liver atlas reveals distinct age-related signatures and provides background against which to interpret pediatric liver disease data.

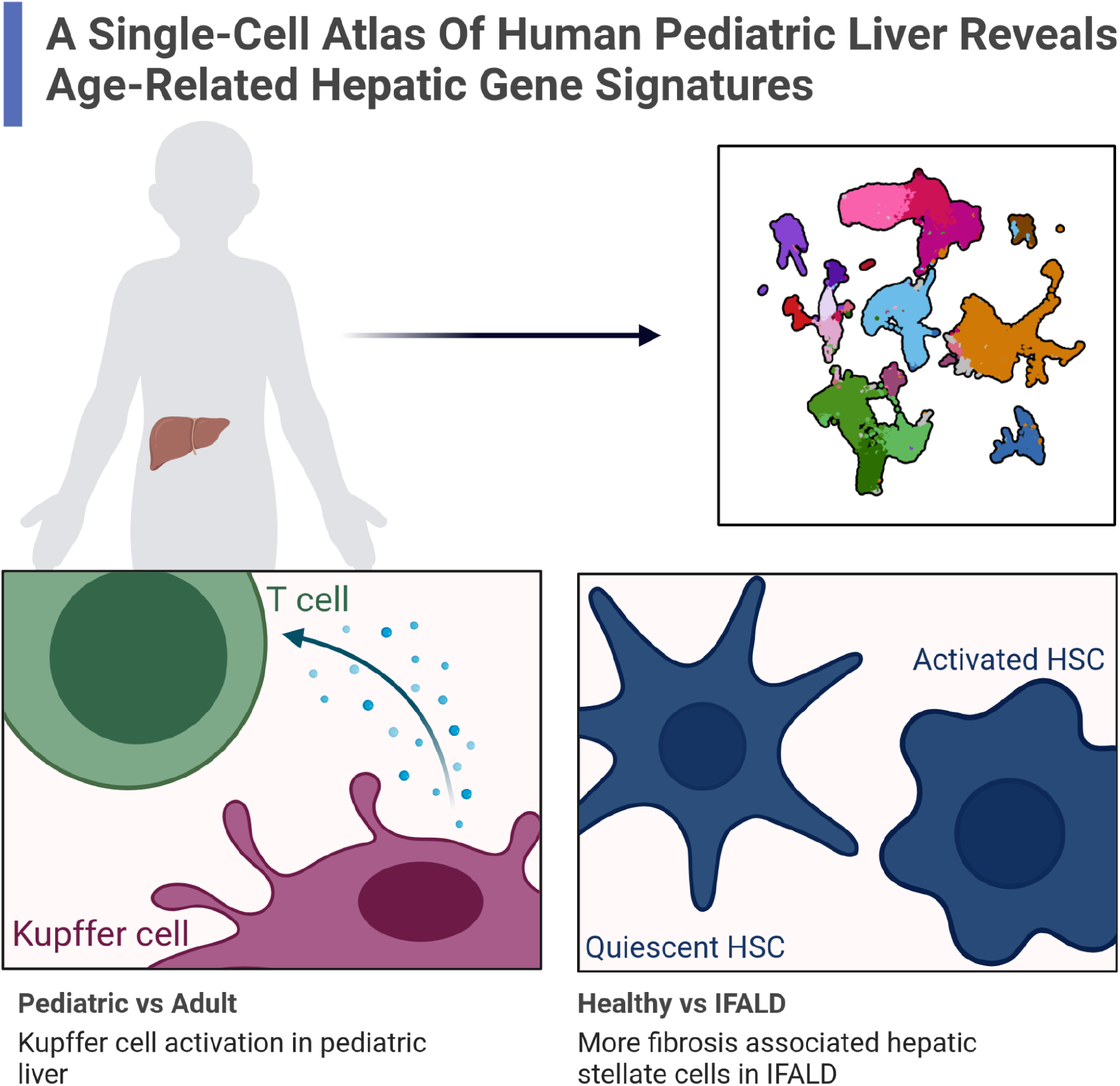

## INTRODUCTION

The liver is vital for human metabolism and immune function. Although the liver is composed of a heterogeneous mix of cell types, how these cell types contribute individually, and in concert, toward liver function remains poorly understood. What is known about healthy liver function, cellular heterogeneity and cell structures is primarily based on studies of the fetal and adult liver ^1–8^. However, there are few studies that have examined the pediatric liver with single cell transcriptomics ^9,10^. The pediatric window is a potentially unique time in liver biology. The body’s immune system is seeded in part from the fetal liver. Subsequently populations of tissue resident immune cells undergo tissue specific maturation programs. How the pediatric liver resident immune system matures and differs from both the fetal liver and fully developed adult liver is unclear.

It is well established that functional differences between pediatric and adult liver exist at the level of metabolism and response to liver damage. For example there are differences in drug metabolism ^11–13^, potentially through differences in cytochrome P450 protein levels ^14^, as well as differences in alanine transaminase levels, which is an indicator of the response to liver damage ^15^. These differences in healthy liver function provide a clear rationale for creating an atlas focused on pediatric livers. Furthermore, there are liver diseases which present differently in children ^16,17^, such as Intestinal Failure Associated Liver Disease (IFALD). When the intestine is unable to absorb adequate nutrition, frequently due to neonatal short bowel syndrome ^18^, parenteral nutrition (PN) is a lifesaving intervention. However, 20%−30% of children receiving prolonged PN will develop IFALD ^19^. Pediatric IFALD is characterized by cholestasis and fibrogenesis while in adults steatohepatitis is seen more commonly^20^. Liver injury seen in IFALD such as cholestasis, inflammation and fibrosis can lead to end-stage liver disease ^21^. The pathogenesis of IFALD is multifactorial, not fully elucidated and is a result of intrinsic and extrinsic factors. More specifically, the intestinal microbiome and components found in PN are expected to play a role in IFALD, with the activation of hepatic macrophages ^22^.

In both the fetal and adult liver, single-cell RNA sequencing (scRNA-seq) has been instrumental in gaining an in-depth understanding of the liver’s cellular heterogeneity and complexity. Based on the relative wealth of adult scRNA-seq data we can now compare adult to pediatric liver and establish commonalities and pediatric-specific differences. A map of the healthy pediatric liver will also enable better understanding of childhood liver diseases, such as IFALD. Here we analyzed healthy human pediatric liver tissue and have made a pediatric liver single-cell map as a community resource. Using this map as a comparator we present the first scRNA-seq map of the normal pediatric liver and the pediatric IFALD liver and identify disease associated differences in cell states.

## METHODS

### Liver Sample Collection

Healthy human liver tissue from the caudate lobe (and in three cases, right lobes) was obtained from neurologically deceased donor livers acceptable for liver transplantation and without evidence of histopathological liver disease some of which has been previously published (**Table S1**). IFALD samples were from children with IFALD undergoing an elective surgery due to their short bowel syndrome and samples are core-needle biopsies. Healthy adult samples were collected with institutional ethics approval from the University Health Network (REB# 14-7425-AE) and healthy and diseased pediatric samples through the Hospital for Sick Children (REB# 1000064039). Samples were collected and processed for scRNA-seq, as previously described ^23,24^.

### Single-cell RNA-seq Data Analyses

Full details are in the Supplementary Material but in brief, sequencing reads were aligned using 10x Genomics Cell Ranger 3.1.0 software to the reference human transcriptome (GRCh38-Ensembl 93; Cell Ranger reference package-3.0.0). The resulting counts were processed using Seurat ^25–28^, SoupX ^29^, *rPCA* ^*30*,*31*^, cell cycle scoring ^32^ and SCINA ^33^. Cell type annotation was based on known cell type markers (**Table S2**) ^1,7,32,34–44,45–51^. Differential expression of individual genes between healthy pediatric and adult or IFALD samples were tested in each cell type, using the *FindMarkers* function in Seurat ^52^. To explore the pathways related to differential expression with age and IFALD, we examined the enrichment of gene ontology (GO) groups fgsea ^53^. Cell-cell interactions were inferred using CellPhoneDB ^54^. To compare to fetal liver ^2^ was done using varimax rotations of the PCs after integration ^55^.

### Spatial Transcriptomics

From 5 livers, 7 FFPE liver sections (one liver sampled three times) were applied to Xenium slides following manufacturers recommended protocols (10x Genomics, Pleasanton, CA, USA) (**Table S3**). then run on the Xenium Analyzer at the Princess Margaret Genome Centre using the Human Multi-Tissue plus 100 liver specific custom markers (**Table S4**). For all spatial data sections used were deemed normal by a pathologist evaluation of H&E stain. One sample (C94_2) contains a bile duct hamartoma so was excluded from analysis. Transcript counts and coordinates were exported from Space Ranger (10x Genomics, Pleasanton, CA, USA). Further details on segmentation using BIDCell ^56^, cell type annotation, and zonation of hepatocytes is in the Supplementary Materials.

### Myeloid Intracellular Cytokine and Immunofluorescent Staining

The IL1-β secretion response by cells was assessed by stimulating biobanked total liver homogenate (TLH) cell suspension specimens with lipopolysaccharide (LPS). Paraffin-embedded sections from healthy pediatric and IFALD liver were stained and quantified as described previously ^57^. Details are provided in the Supplementary Materials and antibodies and listed in **Table S5**.

## Data Availability

The healthy pediatric liver map and IFALD data are available via CELLxGENE. Spatial transcriptomics data are available for exploration through a shiny application macparlandlab.shinyapps.io/pediatric_liver_spatial. Spatial data are available under [*link to be added upon publication*] and the single cell map is in the HCA Data Portal [*link to be added upon publication*] and CELLxGENE: [*link to be added upon publication*].

In the interest of patient privacy pediatric ages are published as a range. Exact age can be provided upon reasonable request. All code for analysis presented here is available at github.com/redgar598/liver_ped_map. AI tools were used to improve readability and language of this work with the authors’ oversight and careful review.

## RESULTS

### Landscape of Pediatric Liver Cells

We generated 54,629 high-quality single-cell liver transcriptomes from 9 pediatric neurologically deceased liver donors (referred to as “healthy”) and 3 IFALD donors (**Table 1 and S1**). For comparison we have used 26,372 liver cells from 7 healthy adult patients (**Table 1 and S1**). Single-cell profiles partitioned into 35 clusters which we annotated using known markers (**Fig. 1A-C, Table S2**). All coarse clusters are well represented across technical variables and generally proportional across individuals (**Fig. S1 and S2;** entropy > 0.48). While distinct parenchymal and non-parenchymal compartments are seen, *ALB* expression was also seen in *PTPRC+* cells suggesting ambient mRNA contamination from hepatocytes in all other cells (**Fig. 1D**). Ambient mRNA was seen to a greater extent in adult samples (**Table S1**), but we expect this is a chance technical effect and not an age related biological effect. All coarse cell types identified in the healthy pediatric map are also seen in adults (except for a population of neutrophils, 70% of which were from one adult sample, ID: C97) (**Fig. 1E-F, S3**).

**Figure 1:**
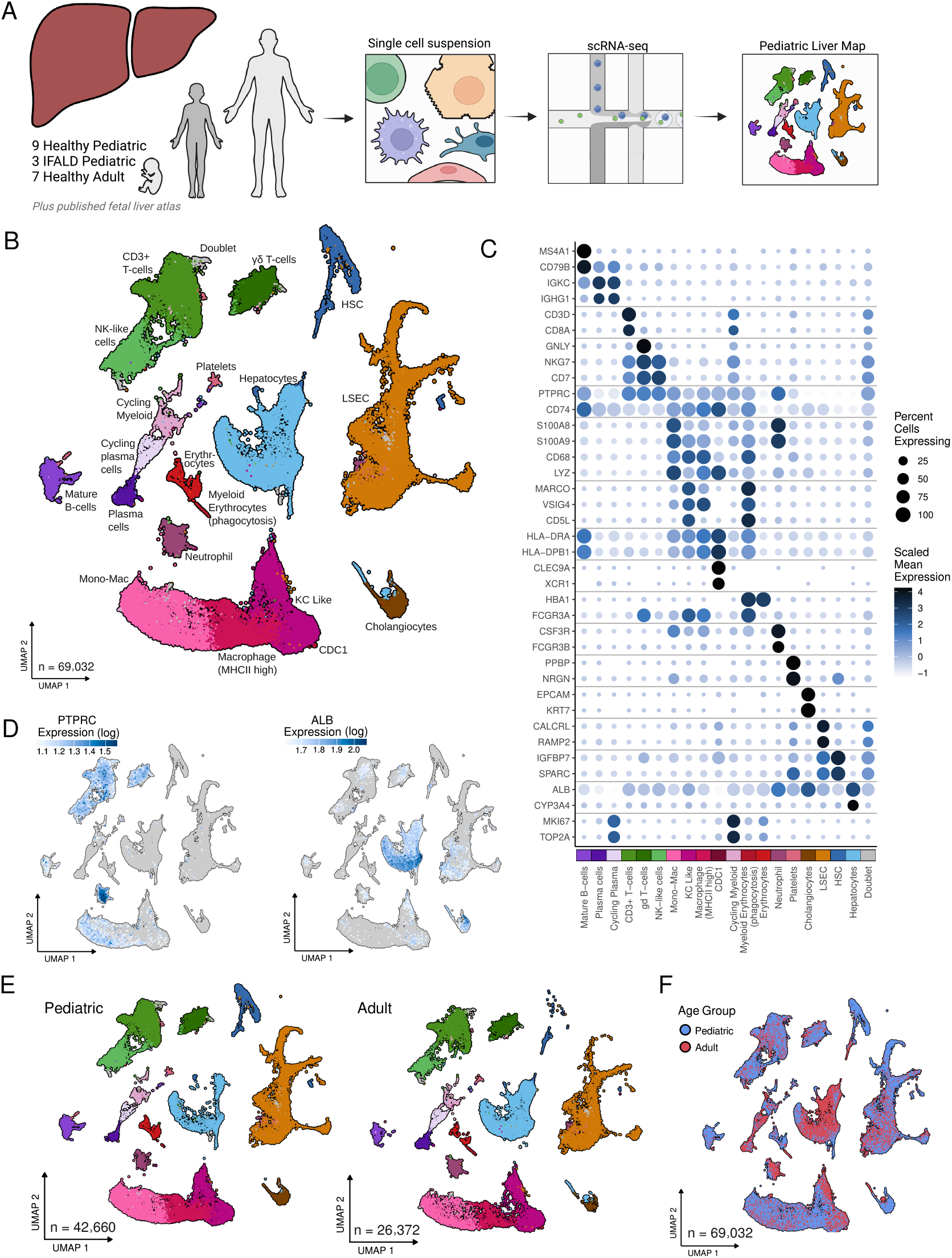
Landscape of cells in the pediatric liver shows similar coarse clusters to the adult liver. A) Overview of map generation process. B) UMAP of combined healthy pediatric and adult liver cells. C) Gene expression of key markers used for cell type annotation. The color of the points represents the gene expression level and size represents percent of cells of a type expressing the gene at all. D) Pediatric liver map colored by expression of a key hepatocyte marker (ALB) and a non-parenchymal cell marker (PTPRC: i.e CD45). E) The pediatric liver map F) split by and F) coloured by age group.

### Immune Activity of Pediatric and Adult Liver Resident Myeloid Cells

In both adult and pediatric livers, myeloid cells selected based on marker genes (**Table S2**) could be sub-clustered into three major populations: Kupffer cell (KC)-like liver resident macrophages (*VSIG4, MARCO* and *CD5L*), recently recruited monocyte-like (Mono-Mac) myeloid cells (*S100A8/9* and *LYZ*) or major histocompatibility class II (MHC II) high myeloid cells (*HLA-DRA/DPB1/DQB1*) (**Fig. 1B and 2A**). A minor population of MHC II and *CLEC9A* high cells was seen in both pediatric and adult livers. These may be conventional type 1 dendritic cells (cDC1) ^58^, but as there were only 38 of these cells in the healthy pediatric and adult samples, little can be concluded from this population.

When comparing pediatric to adult in each of the three major myeloid populations, genes associated with immune activation were more highly expressed in pediatric livers. In KC-like cells in particular, key chemokines (*CCL3* and *CCL4*) were more highly expressed as well as *IL1B* (**Fig. 2B and Table S6**). The immune activation gene signature seen in pediatric KC-like cells has previously been seen in fetal liver macrophages ^2,59^. Combined fetal and pediatric liver data we saw even higher immune activation in the fetal liver (**Fig. S4**, further detail in Supplementary Materials). Suggesting the KC-like immune activation may be highest in fetal liver and wane through gestation and aging. These genes and others contributed to a significant enrichment of the “*IL-18 signaling*” pathway in genes more highly expressed in pediatric KC-like cells (**Fig. 2B and S5**). In the MHC II high myeloid cells *HMOX1* and *APOE* are more highly expressed in pediatric cells and drive an enrichment of the pathway “*regulation of epithelial cell proliferation*” (**Fig. 2B and S4; Table S7**). In the Mono-Mac cells, genes associated with cellular stress pathways were more expressed in pediatric cells with *AREG* ^60^ and several heat shock protein-related genes were more highly expressed (**Fig. 2B and S5; Table S8**).

**Figure 2:**
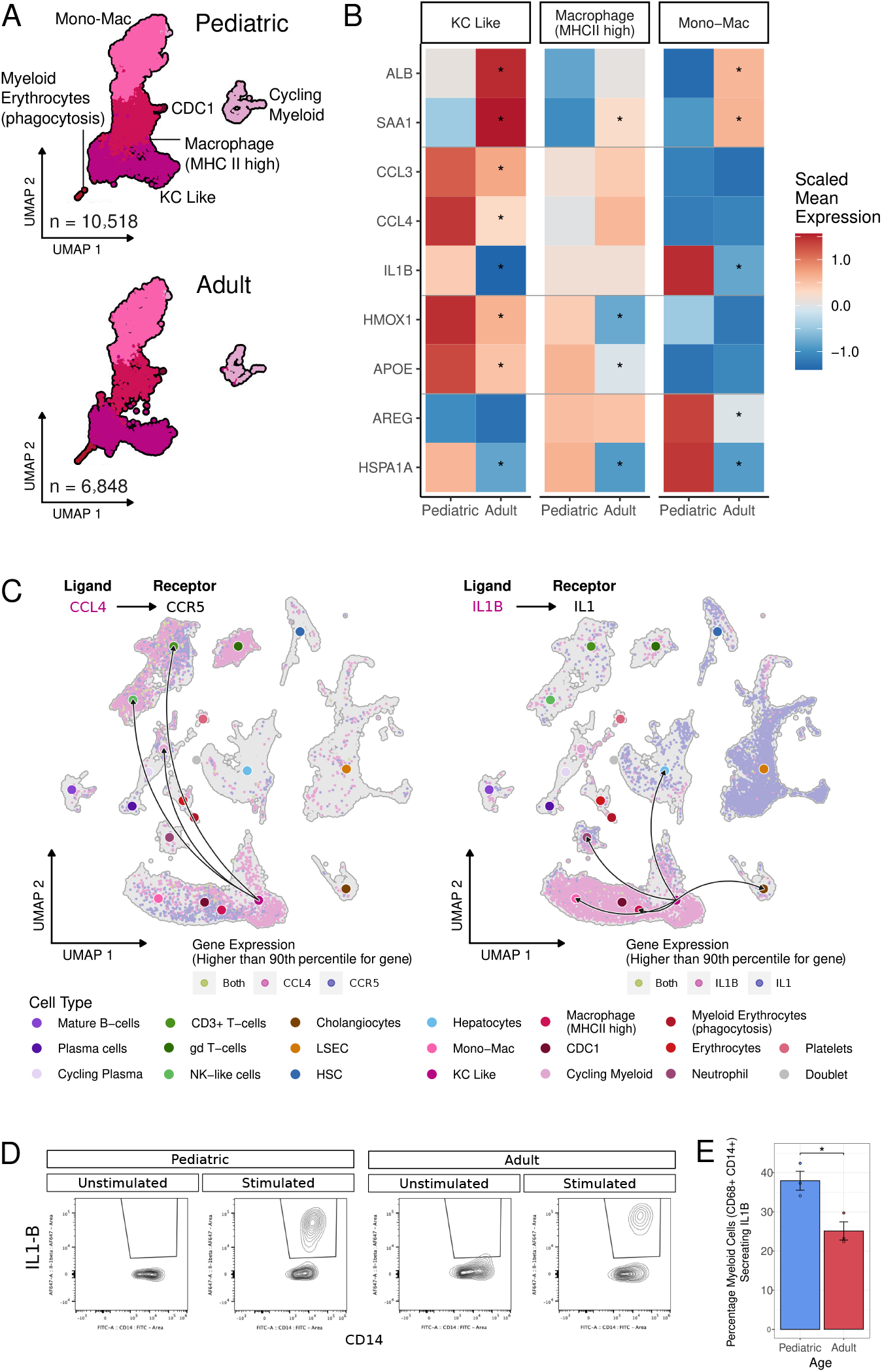
Pediatric liver myeloid cells express more inflammatory cytokines than adult cells. A) UMAPs of liver myeloid cells split by sample age group. B) Differentially expressed genes in each myeloid cell type. Rows show individual genes and columns show samples split by age group. Horizontal lines divide genes into following categories: higher expression in adults in all myeloid types, higher expression in pediatric KC-like, higher in pediatric in MHCII high cells and higher expression in pediatric in mono-mac cells. C) Predicted cell-cell interactions between KC-like cells and all other cell types. Individual cells in the UMAP are coloured by the expression of the ligand and/or receptor. Left panel: CCL4-CCR5 right panel: IL1B-IL1. Arrows on the connecting curves point from ligand to receptor and curves are only shown for significant cell-cell interactions. Larger points which curves connect are colored to indicate the cell population. D) Percentage of IL1-β^+^ secreting myeloid cells (CD68^+^CD14^+^) in the unstimulated control and LPS stimulated conditions of pediatric and adult myeloid cells. E) Stimulated adult versus pediatric cells with percent of myeloid cells (CD68^+^CD14^+^) secreting IL1-β shown.

Focusing on the KC-like cell immune activation, we observed that pediatric KC-like cells are predicted to interact with other myeloid cells and T cells *via* known ligand-receptor pairs. Specifically *CCL3* and *CCL4* expressed in KC-like cells could be interacting with Mono-Mac and MHC II high myeloid cells (through *CCR1*) and T cell populations (through *CCR5* and *CCR1*) (**Fig. 2C**), suggesting KC in the pediatric liver could be recruiting T cells and promoting monocyte chemotaxis. Looking at interactions inferred from expression of known IL1-β binding partners, interactions were predicted between KC-like cells and many other cell types: hepatocytes, cholangiocytes, neutrophils, Mono-Mac cells and MHC II high myeloid cells (**Fig. 2C**). However, there were also predicted interactions between IL1-β and the IL1-β receptor inhibitor in neutrophils, Mono-Mac cells and MHC II high myeloid cells, but not in hepatocytes and cholangiocytes (**Fig. S6**). So potentially the interaction of IL1-β with its receptor is inhibited in myeloid cells but not in hepatocytes and cholangiocytes allowing activated pediatric KC-like cells to signal to parenchymal and epithelial cells specifically.

From these data, we asked the question of whether the higher expression of immune related genes could be linked to protein level functional differences related to cytokine and chemokine secretion. Having seen *IL1B* more highly expressed in the pediatric liver we tested the capacity of myeloid cells (CD68^+^CD14^+^) isolated from pediatric and adult livers to secrete IL-1β upon LPS stimulation. We found a significantly higher frequency of pediatric myeloid cells secreting IL-1β in response to LPS compared to adult myeloid cells (p<0.05; **Fig. 2D-E**). This confirms the immune activation signature seen in the pediatric scRNAseq map does confer a higher inflammatory potential compared to the adult liver.

Zonation of immune cells within the liver is a dynamic system. We therefore examined the zonation of gene expression in myeloid cells genes to understand if spatial dynamics in liver transcriptomics could explain some differences seen between the pediatric and adult liver. We explored the zonation of a panel of 477 genes using the Xenium spatial transcriptomics platform in two pediatric and three adult livers (**Table S2**; **Fig. 3A-C, S7**). Of the 70 genes significantly differentially expressed in pediatric KC-like cells in the single-cell pediatric map, 10 genes were included in the spatial panel. Of these, 8 genes were also zonated in KC-like cells (FDR< 0.005; **Table S9, Fig. 3D-F**). This included KC marker genes (*MARCO* and *CD5L*) which were zonated within KC-like cells in all samples (**Fig. 3F**). The zonation of the KC markers genes suggests the observed differential gene expression in the dissociated map could be the result of underlying shifts in KC zonation in the pediatric liver.

**Figure 3:**
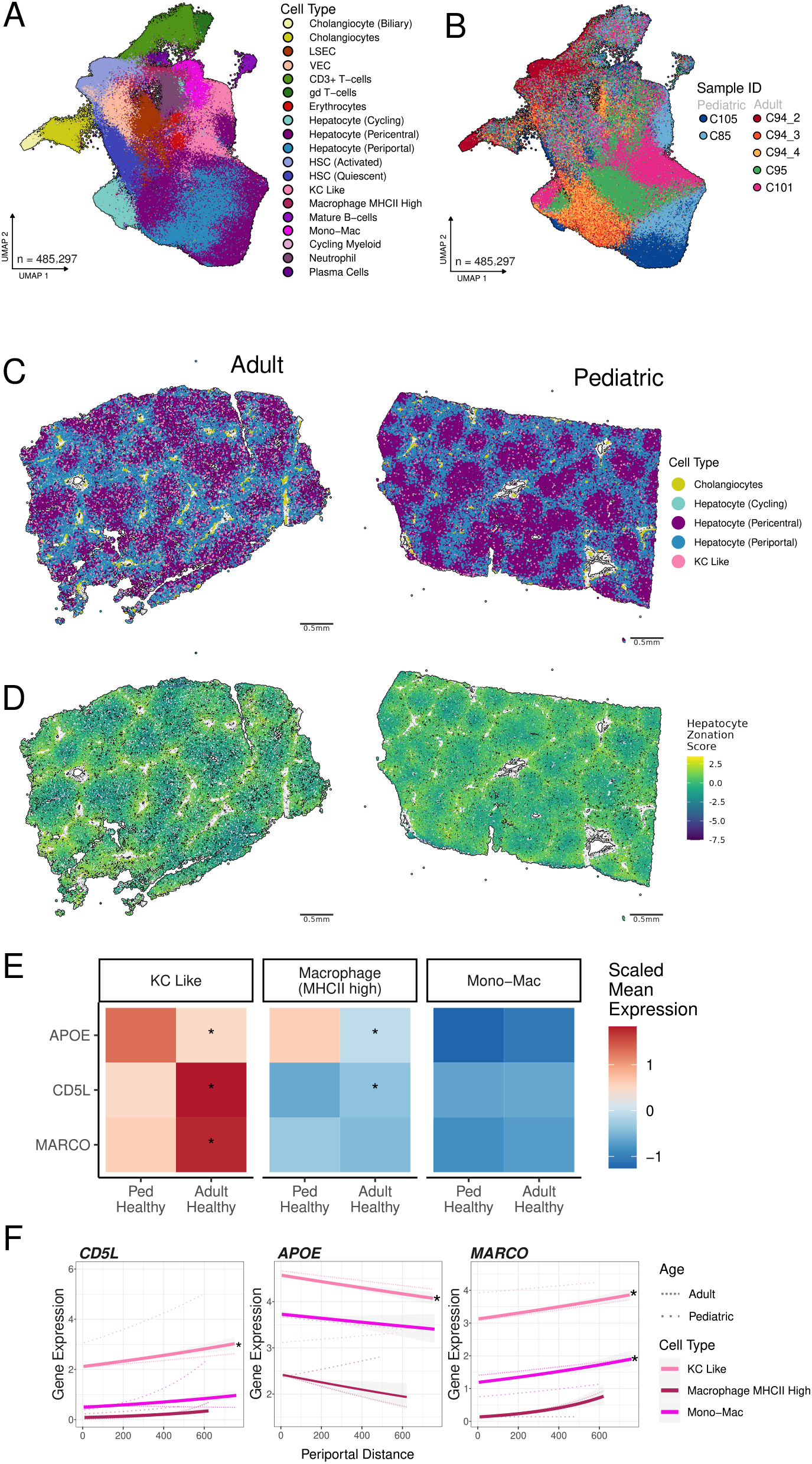
Zonation is successfully captured in both pediatric and adult spatial transcriptomics at the single cell level. UMAP of combined healthy pediatric and adult liver cells from the spatial transcriptomics assay coloured by A) cell type and B) donor ID. C) Representative adult (C94_4) and pediatric (C105) spatial transcriptomics samples with points representing cells coloured by cell type and positioned in their physical location in the tissue. Only the listed cell types are shown to highlight the zonation between portal and central veins. D) Only hepatocytes are shown with cells coloured by zonation score. E) Differential expression, in the dissociated single-cell map, of genes present in the spatial data. F) The same genes are significantly differentially expressed with inferred distance to a portal vein in KC-like cells in the spatial transcriptomics samples. Thicker lines represent the trend across samples with thinner lines for the trend in pediatric and adult separately.

Of note one sample from C94 (C94_2) captured a bile duct hamartoma (**Fig. S8**). This sample was excluded from analyses but the data will be released along with the other spatial data presented here as it may be of interest to the community.

### Fibrosis and Immune Activity in IFALD Liver

Having established the healthy pediatric map, we then applied this map as a comparator to explore the cellular complexity of liver biopsies from pediatric IFALD patients (**Fig. 4A**). In the combined adult, healthy pediatric and IFALD map we observed two distinct populations of hepatic stellate cells (HSCs) (**Fig. 4B**). HSC population one is made up of mainly healthy pediatric HSCs (92% of cells) and HSC population two is made up of mainly IFALD (48% of cells) and adult HSCs (23% of cells), suggesting HSCs in IFALD are more similar to healthy adult than healthy pediatric HSCs. Comparing these two HSC populations we saw fibrosis genes (*PDGFRA, CXCL12, COL1A1* and *IGFBP3*) more highly expressed in HSC population two which is mainly adult and IFALD cells (**Fig. 4C; Table S10**). Fibrosis is a key feature of IFALD and it is interesting to confirm HSC activation at the single-cell level. Without the healthy pediatric map, IFALD HSCs would have only been compared to healthy adults and this evidence of fibrosis at the single-cell level may have been missed as several fibrosis markers (*PDGFRA, CXCL12*, and *IGFBP3*) were not differentially expressed when comparing IFALD to adult HSCs (**Table S11**).

**Figure 4:**
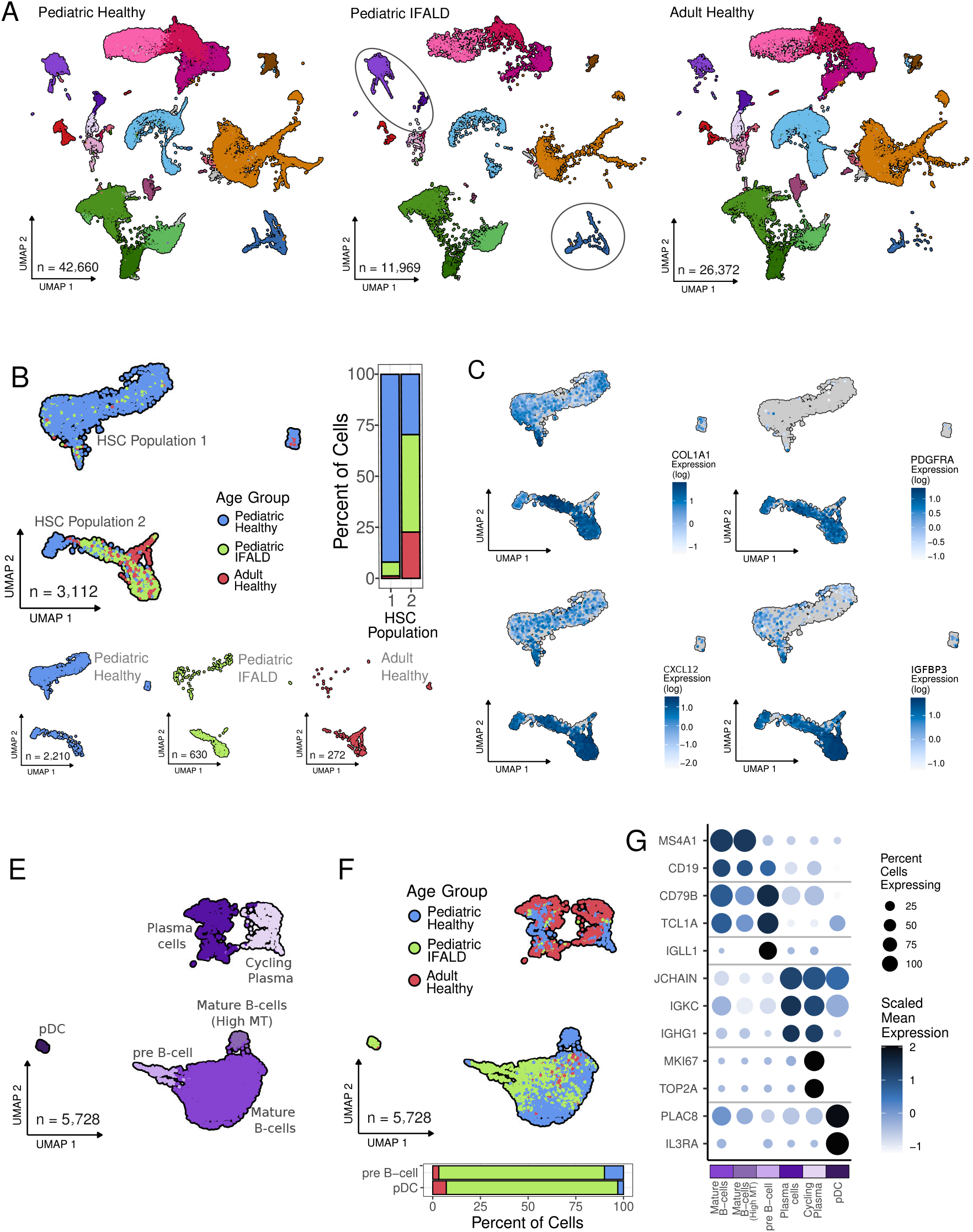
IFALD HSCs are more fibrotic than non-diseased livers and contain unique B cell types. A) UMAP of liver cells with key cell types highlighted where differences in IFALD are described. B) HSC UMAP coloured as well as split by age and disease status, with a barplot showing the composition of each HSC population. C) UMAPs colored by differentially expressed fibrosis genes. UMAP of B cells coloured by E) cell type or F) sample age and disease group, with a barplot show the composition of pre-B cell and pDCs. G) Gene expression of key B cell markers used for B cell annotation. The color of the points represents the gene expression level and size represents percent of cells of a type expressing the gene at all.

The IFALD HSC findings highlight the limitations of using healthy adults as a pediatric disease comparator, an alternate approach used previously has been employing other pediatric liver diseases as comparators ^61^. Looking at cholangiocytes from a previously published Biliary Atresia map where the original comparator was cells from patients Choledochal Cysts, we observed higher expression and pathway enrichment of MHCI and MHCII signaling in both Biliary Atresia and Choledochal Cysts compared to healthy pediatric liver (See Supplementary Material; **Fig. S10 and S11**). As this signal is shared in cholangiocytes from patients with Biliary Atresia and Choledochal Cysts it was not observed in the original publication.

Turning to immune populations, we saw several populations unique to IFALD and not seen in either healthy pediatric or adults. Specifically, we saw more pre-B cells (*IGLL1*+ but *MS4A1*-)^62,63^ in IFALD. We also identified plasmacytoid dendritic cells (pDCs; *IL3RA+* and *PLAC8+*) ^2^ which were more abundant in IFALD (**Fig. 4E-G**). We were also interested in any macrophage specific differences in IFALD. In terms of differences in cell type abundance, we see similar levels of major cell types (**Fig. 5A**). We did see the cDC1 population more abundant IFALD (**Fig. 5A**) and cDCs have been seen previously to be more abundant in metabolic dysfunction-associated steatohepatitis (MASH) ^64^. However, both the pDC and cDC1 differences could result from the differences in sample processing (See Supplementary Material; **Fig. S9AB**).

**Figure 5:**
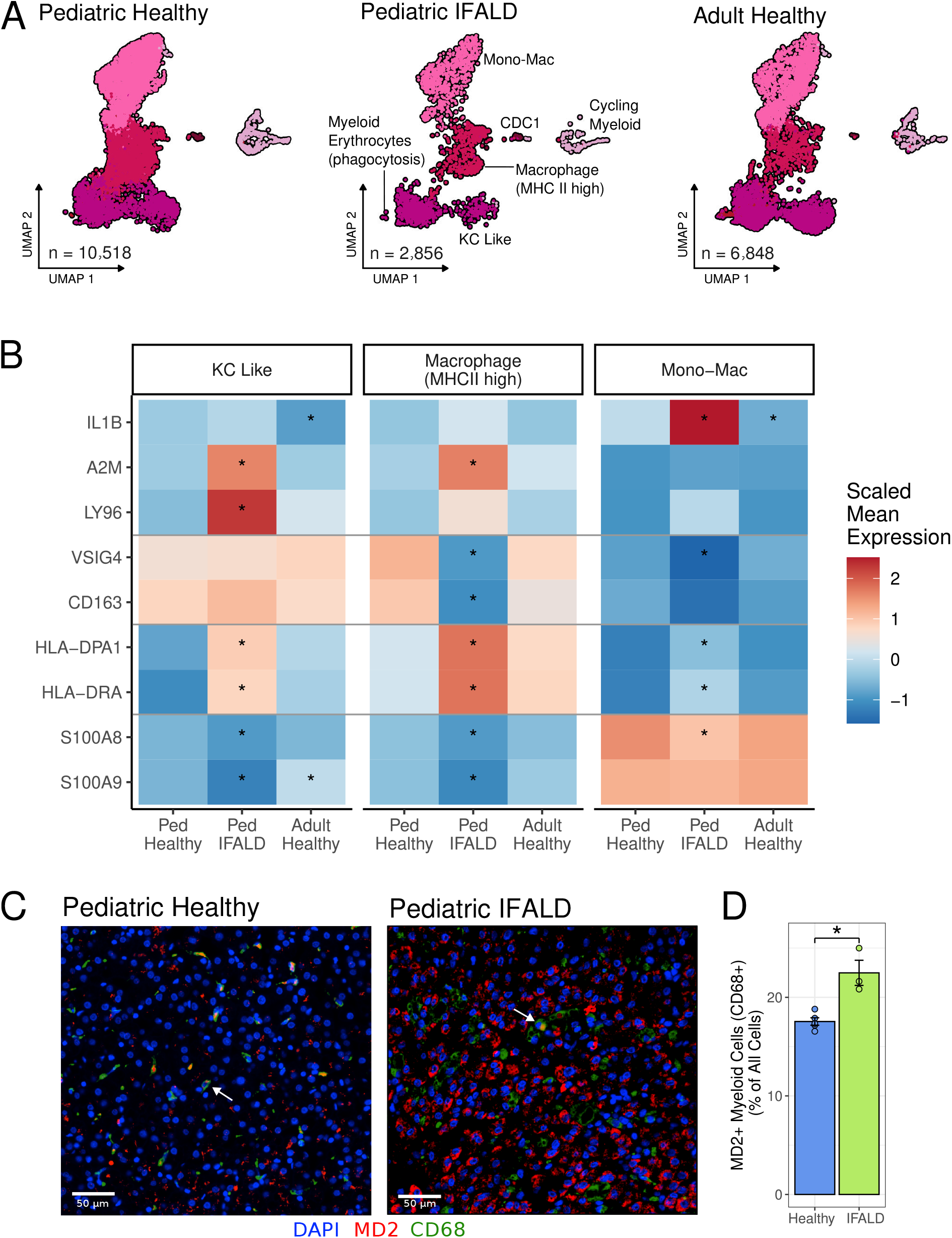
IFALD myeloid cells more highly express profibrotic genes than non-diseased livers. A) UMAP of liver myeloid cells split by age and disease. B) Differentially expressed genes for each myeloid cell type. Rows show individual genes and columns show samples split by age and disease. The horizontal lines divide genes into the following categories: higher expression in IFALD in KC cells, lower expression in IFALD MHCII high cells, higher expression in IFALD MHCII high cells, and lower expression in IFALD KC like and MHCII high cells. C) Representative IF images of pediatric healthy and IFALD livers. D) Quantification of the MD2^+^CD68^+^ cells.

When comparing healthy pediatric to IFALD in the three major myeloid populations, we saw potential shifts in cell identity and function. In Mono-Mac myeloid cells we see *IL1B* more highly expressed in IFALD than in healthy pediatric (**Fig. 5B and Table S12**), suggesting a similar immune activation seen when comparing pediatric to adult. In addition we observed a potential profibrotic state of IFALD myeloid cells. In KC-like cells we saw higher expression of profibrotic genes (i.e *LY96*) (**Fig. 5B and Table S13**) ^65^. The overexpression of *LY96* unique to IFALD and not seen in comparison to other pediatric liver diseases (See Supplementary Material; **Fig. S12**). As the protein product of *LY96* (MD-2) is a component of Toll-like receptor 4 (TLR4) ^66^ we wanted to validate its increased abundance in IFALD myeloid cells at the protein level. Using immunofluorescence staining of healthy pediatric and IFALD samples we confirmed the higher proportion of MD-2^+^CD68^+^ myeloid cells in IFALD compared to healthy pediatric myeloid cells (p<0.05; **Fig. 5CD**).

## DISCUSSION

The pediatric liver represents an important but understudied window of development. Using single-cell transcriptome profiling we describe the cellular microenvironment of the pediatric liver in comparison to adults and pediatric liver disease. This map is a resource for the community to study pediatric liver disease. Inherent differences, like the immune activation in pediatric myeloid cells, mean the pediatric map will serve as an important comparator to account for normal differences with age in the liver, as well as with disease conditions.

### The function of macrophage immune activation during liver development is unclear but a similar trend has been seen in other age ranges and tissues

In fetal tissues, including liver, there is greater macrophage immune activation at earlier gestational ages ^59^. A possible function for the immune activation is suggested by the higher expression of genes related to epithelial cell proliferation in MHC II myeloid cells (*HMOX1* and *APOE*) and the potential for myeloid cells to be signaling cholangiocytes and hepatocytes via *IL1B*. Shifts in liver immune composition have been seen to be a crucial element in healthy liver development ^67^ The observed immune activation of KC-like cells could be a normal part of the growth and structural development of parenchymal and epithelial cells in the pediatric liver, a process which is later suppressed in adulthood, once the liver is fully formed.

### Characterizing the pediatric liver could have important clinical implications

Age-related differences in the chemokines we have seen in the liver are also seen in pediatric blood compared to adult blood ^68^. There is an observed decline with age in the abundance of the proteins CCL4, IL-1β and CXCL8 over a similar age range as our samples, but no association between CCL3 and age was seen ^68^. In a separate cohort, CCL4 and CXCL8 abundance in blood also declined with age, but again CCL3 was not significant ^69^. Potentially, the immune activation we have seen in the pediatric liver is involved in shaping the liver’s growth and structure, and the age-related differences in these secreted chemokines can be seen in blood.

### The differences we have found between pediatric and adult livers illustrates the value of an age matched comparator for studying pediatric liver disease

The signal (*COL1A1*+, *PDGFRA*+, *CXCL12*+ and *IGFBP3*+) of fibrosis in IFALD-enriched HSCs was expected given that IFALD is known to involve liver fibrosis, ^45,47,48,70^. However if IFALD HSCs had only been compared to adult HSCs then this fibrosis would likely have been missed. Fibrosis, observed at the single-cell level in IFALD, is an important observation as it is similar to the signal seen in MASH and may suggest some pathogenic similarities between the two diseases ^65,71^, but is distinct from differences seen in single-cell transcriptomics of primary sclerosing cholangitis (PSC) ^8^. As Metabolic dysfunction-associated fatty liver disease (MAFLD), MASH and PSC all present uniquely in children compared to adults the finding of immune activation in the pediatric liver may explain some of the clinically observed differences if the pediatric liver is generally a more immune active background for the disease ^72,73^. These differences in immune activation may also explain the different histology findings in IFALD between children and adults with the disease. In addition to the fibrosis signature seen in HSCs, differences in myeloid subpopulations would likely also have been missed if pediatric IFALD was only compared to healthy adults. We saw *IL1B* more highly expressed in IFALD Mono-Mac cells, suggesting an IL-18 signaling role in IFALD in Mono-Mac cells. We did not see the same higher expression of *IL1B* in IFALD KC-like cells compared to healthy pediatric livers, but *IL1B* in IFALD KC-like was different from adult KC-like cells, likely due to the generally higher immune activation of the pediatric liver. Our findings support previous publications showing the major role of IL-1β in the pathogenesis of IFALD through down regulation of LXR and canalicular ABCG5/G8 ^19^. As with HSCs, if we had only compared IFALD to healthy adults the immune activated state of the pediatric KC-like population may have been incorrectly attributed to IFALD.

### With this age matched comparator we were able to observe a potential role of myeloid cells in contributing to IFALD fibrosis

We have seen lower expression of *S100A8* and *S100A9* in IFALD myeloid cells, and this is similarly seen in metabolic dysfunction-associated fatty liver disease (MAFLD) myeloid cells ^74^. As well a similar profibrotic (i.e. *LY96*+) role of IFALD KC-like cells is also seen in MASH ^65^. One potential mechanism in the pathogenesis of IFALD is the crossing of lipopolysaccharides (LPS) from the gut into the portal system due to leaky gut, typical of children with short bowel syndrome ^22^. The protein encoded by *LY96*, MD-2, is a component of TLR4, the core receptor for LPS. The *LY96*+ KC like population in IFALD may support the theory that inflammation and eventual failure of the liver in part occurs through LPS driven inflammation at portal veins. This is a promising finding as MD-2 is a druggable target with inhibitors being developed for the treatment of systemic sclerosis ^75^.

### The healthy pediatric and IFALD maps will be improved with the continual integration of more samples

Some technical confounders can not be fully controlled for in the current analysis, such as the adult samples used for comparison having more ambient mRNA contamination and the IFALD samples being from liver biopsies versus perfused caudate lobes from neurologically deceased donors used for the healthy pediatric map. These can be overcome with future inclusion of more diverse samples, integrated across labs ^76^. Another confounder between pediatric IFALD and the healthy pediatric samples, which may explain some of the cell populations unique to IFALD, is a difference in age. Two of the three IFALD sample donors were under one year of age and the youngest healthy donor was a two year old patient. As the liver is a site of hematopoiesis during gestation, the abundance of pre-B cells in IFALD may be a normal feature of livers under one year of age and not a unique feature of IFALD. This is supported by the observation of similar pre-B cells in the fetal liver ^59^. As pediatric liver samples are exceedingly rare it is important to share this map even with the current caveats. This resource has enabled key insight on pediatric liver disease and will be a valuable resource to the community.

## List of Abbreviations

AI: Artificial Intelligence
ALT: Alanine Transaminase
cDC1: Conventional Type 1 Dendritic Cells
DC: Dendritic Cells
DAPI: 4’,6-diamidino-2-phenylindole
FDR: False Discovery Rate
FFPE: Formalin-Fixed Paraffin-Embedded
FMO: Fluorescence Minus One
FGSEA: Fast Gene Set Enrichment Analysis
GO: Gene Ontology
GRCh38: Genome Reference Consortium Human Build 38
H&E: Hematoxylin and Eosin
HCA: Human Cell Atlas
HSC: Hepatic Stellate Cells
IFALD: Intestinal Failure-Associated Liver Disease
KC-like cells: Kupffer Cell-like cells
LPS: Lipopolysaccharide
MAFLD: Metabolic Dysfunction-Associated Fatty Liver Disease
MASH: Metabolic Dysfunction-Associated Steatohepatitis
Mono-Mac: Monocyte-like Macrophages
NPC: Non-parenchymal Cells
PBMC: Peripheral Blood Mononuclear Cells
PC: Principal Component
PCA: Principal Component Analysis
PN: Parenteral Nutrition
PSC: Primary Sclerosing Cholangitis
REB: Research Ethics Board
rPCA: Robust Principal Component Analysis
scRNA-seq: Single-cell RNA sequencing
TLH: Total Liver Homogenate

## ACKNOWLEDGEMENTS

This publication is part of the Human Cell Atlas (humancellatlas.org/publications). The authors acknowledge the Princess Margaret Genomics Centre, the Pathology Research Program and the Advanced Optical Microscopy Facility at University Health Network for their support and services. The graphical abstract was created with Biorender.com. The authors would like to acknowledge Dr. Michael Cheng and Dr. Harry Sutton for comments and insights on the study. The authors acknowledge Dr. Stacey Huppert for informal peer review of this manuscript.

