## Supplementary Material for "A Single-Cell Atlas Of Human Pediatric Liver Reveals Age-Related Hepatic Gene Signatures"

### Supplementary Methods

#### Single-Cell Quality Control

Counts were adjusted for estimates of the ambient RNA in a cell using SoupX[29]. Cells with less than 500 or more than 6,000 detected genes and cells with mitochondrial transcript proportion higher than 25% were excluded. Data were normalized using sctransform in Seurat[30,31] and the cell cycle timing [32] and number of genes detectable in a cell (nFeature_RNA) were regressed out at this step.

#### Single-Cell Integration

Data was integrated across patients (healthy pediatric, adult and IFALD) using the Seurat function rPCA, with anchors found from the 3,000 most variable genes[25–28]. The resulting integrated gene expression counts were used for visualization, clustering and cell type annotation but non-integrated, log normalized counts were used for quantitative comparisons. Separately, but using the same procedure, liver samples were integrated with the one PBMC sample.

#### Single-Cell Differential Expression

Cells were clustered using the Seurat functions FindNeighbors and FindClusters (30 PCs and a resolution of 0.5). Then clusters were assigned a cell type label based on SCINA 33 labeling using known cell type markers (Table S2) [1,7,32,34–44]. Labels were then confirmed and/or refined with sub-clustering and manual annotation labels with known cell type markers (Table S2; "Subtype") [45–51]. Clusters of cells expressing multiple markers (often immune cells expressing LSEC markers) were annotated as doublets, except for cells expressing both myeloid and erythrocyte markers, these were labeled "Myeloid Erythrocytes (phagocytosis)".

To check that cells in clusters were well mixed after integration across technical factors (individual, chemistry, age and disease groups), Shannon entropy was calculated for the mixing of factors in each of the 32 clusters. We set a threshold for low entropy based on a theoretical cluster composed of 95% of cells from one individual/age group etc. and the remaining 5% of cells distributed to all other individuals/age groups [52]. Clusters were considered low entropy below the thresholds of 0.49, 0.34, and 0.29, for individual, age and disease condition, and assay chemistry, respectively.

For differential gene expression, the counts were log-transformed and scaled by the total counts in a given cell and multiplied by 10,000. Differential expression of individual genes between healthy pediatric and adult or IFALD samples were tested in each cell type, using the FindMarkers function in Seurat, with MAST used for the actual testing and number of genes detectable in a cell (nFeature_RNA) and sex as latent variables [53]. Differential expression was only tested for genes that were detected in more than 10% of cells in any age/disease group. Unless otherwise indicated, all reported significantly differentially expressed genes have an absolute log2 fold change greater than one and a Bonferroni adjusted p value from the MAST likelihood-ratio-test less than 0.005.

To explore the pathways related to differential expression with age and IFALD, we examined the enrichment of gene ontology (GO) groups (Biological Processes only with Inferred from Electronic Annotation pulled Oct. 26 2022) using fgsea [54] with 5% FDR and 10,000 permutations.

Cell-cell interactions were inferred between KC-like cells and all other cell types in healthy pediatric samples only using CellPhoneDB [55]. Significant interactions between cell types, based on the CellPhoneDB list of interacting pairs, were predicted and ligand or receptors differentially expressed between healthy pediatric and adult KC-like cells were highlighted.

#### Spatial Transcriptomics

These quantified transcripts along with the DAPI nuclear staining were used for cell boundary segmentation with BIDCell [57]. For segmentation all parameters can be found in the provided code, but in brief the cell type specificity of the Xenium panel genes was calculated using the dissociated pediatric map data. Cells from the Xenium were then integrated with the pediatric map on the shared genes using rPCA from Seurat [25–27]. Cell annotations were transferred to Xenium cells based on Louvain clustering of the integrated data clustering [25–27]. These transferred annotations were confirmed with the marker info from the panel (Fig. S6). In the cases where no annotated cell type was clear, these clusters were visualized spatially and a physical region label was given to the cluster. Zonation of hepatocytes was captured by PC2 from a PCA of all hepatocytes. This association was confirmed with known zonation markers (CYP1A2 and CYP2A6) and directionality (periportal vs pericentral) based on cholangiocytes at portal veins (Fig. S6). To infer the zone of non-hepatocytes local maxima and minima of the zonation PC were identified. Distance threshold and k neighbors were set for each sample separately to obtain 1 maxima or minima per 100 hepatocytes. Differentially gene expression was then done with distance from the nearest maxima or minima in each cell type.

#### Myeloid Intracellular Cytokine Staining

The IL1-β secretion response by cells was assessed by stimulating biobanked total liver homogenate (TLH) cell suspension specimens with lipopolysaccharide (LPS). TLH from both healthy caudate samples and pediatric samples were plated in 12-well plates for 4 hours, followed by a wash and stimulation with 75ng/mL of LPS for 20 hours. For the last 6 hours, Monensin and Brefeldin were added at a 1:1000 concentration to trap IL1-β inside intracellular compartmentsLive myeloid cells were stained and identified as Live/Dead Aqua (BioLegend) negative, CD3- (Thermofisher, 414-0032-80), CD45+ (BioLegend, 304824), CD68+(BioLegend, 333821), CD14+(BioLegend, 325603) cells (Supplementary Table S3). IL1-β (Thermofisher, 51-7018-42) was quantified by flow cytometry using a SONY ID7000 spectral flow cytometry analyzer. Gating for cell surface and intracellular markers was established using Fluorescence Minus One (FMO) controls for each marker, with IL1-β gating further referenced against fluorescence levels in both FMO and unstimulated controls FlowJo was used to analyze all events. An unpaired t-test was used to evaluate the means between two groups.

#### Immunofluorescent Staining

Paraffin-embedded sections from healthy pediatric and IFALD liver were stained by the Pathology Research Program (PRP) at the Toronto General Hospital according to standard histological procedures. Paraffin-embedded tissues were stained with antibodies for CD68 (Agilent, PG-M1), MD-2 (Abcam, ab24182) and DAPI (Table S5). The stained slides were scanned by the University Health Network Advanced Optical Microscopy Facility using a Leica Aperio AT2 whole slide scanner (Leica Microsystems, Carlsbad CA), and converted into digital images. QuPath software version 0.4.3 software was used for quantification of stain positive cells on the images [58]. A Wilcoxon rank-sum test was used to evaluate means between two groups.

##### Public Data Integration

Cell Ranger outputs were downloaded from two previously published maps (GSE176189 and GSE163650)[9,61]. As with the healthy map, samples were integrated with rPCA. Then cell type labels were transferred from the healthy pediatric map to the public data based on clustering of the combined healthy and diseased maps. To replicate the analysis presented previously[61] differential expression was validated at an adjusted p value $<0.05$ and a fold change $>1.5$ for the comparisons in cholangiocytes. To validate IFALD differential expression in KC like cells the levels used in the initial comparison of IFALD to healthy pediatric were used with the other diseases log2 fold change $>1$ and adjusted p value $<0.005$.

#### Supplementary Results

##### Fetal and Pediatric Liver Comparison

The immune activation gene signature seen in pediatric KC-like cells has previously been seen in fetal liver macrophages [59]. Namely the expression of *CCL3*, *CCL4* and *IL1B* has been seen to decrease in fetal liver macrophages with increasing gestational age. To compare the expression of genes associated with immune activation from fetal, pediatric and adult we integrated our map with previously published fetal liver scRNAseq data [59]. Due to technical differences (i.e. fetal livers were not perfused) and biological differences (i.e. the liver is a site of hematopoiesis during development) there are substantially more erythrocytes captured in the fetal samples. We saw higher expression of erythrocyte genes in fetal KC-like cells, suggesting a contamination of KC-like cells with erythrocyte mRNA. To separate the signature of erythrocyte genes in the fetal KC-like cells from any immune activation signatures we used varimax principal component analysis (PCA) [55] to identify an independent immune activation signature. The second varimax PC was enriched for the same immune activation genes (*CCL3*, *CCL4* and *IL1B*) seen in the differential expression analysis. We saw that PC2 was significantly higher in fetal and pediatric samples compared to adults (Fig. S6), suggesting the KC-like immune activation is highest in fetal liver and wanes through gestation and aging.

##### Immune Activity in IFALD Liver

The majority of healthy pediatric samples were from caudates which were perfused and dissociated to single-cells, with two healthy pediatric samples from liver biopsies. The IFALD samples were all biopsies (Table 1). Biopsies, by nature, are not perfused and likely include more peripheral blood cells than perfused caudate samples. Possibly peripheral blood in the IFALD biopsies contributed to the observed greater abundance of pre-B cells and pDC in IFALD. We examined this confounding in a separate analysis where we integrated one pediatric peripheral blood mononuclear cell (PBMC) sample with the liver map, to understand if the cell populations unique to IFALD overlap with PBMC populations (Fig. S9AB). Here we saw that dendritic cells (DC) from PBMCs and the liver pDC population clustered together in the integrated PBMC and liver map (Fig. S9CD). Given overlapping transcriptomic profiles of PBMC DC and liver pDC we can not be confident the IFALD pDC was a true liver population. The pre-B cell population however is distinct from the PBMC cell types and is potentially a true liver resident population which was more abundant in IFALD (Fig. S9CD). There is overlap in the there is overlap between the liver cD1 and PBMC cDC populations as well, making it difficult to conclude whether the cDC1s are actually more abundant in IFALD.

We saw possible shifts in myeloid cell identities in IFALD toward an MHC II high signature. We observed all IFALD myeloid cells more highly expressed MHC II genes compared to healthy pediatric (Fig. 5B) and the MHC II high macrophage population expressed comparatively less KC-like and Mono-Mac myeloid markers (*VSIG4* and *S100A8/9*, respectively) in IFALD than healthy pediatric cells (Fig. 5B and Table S14).

#### Comparison to Existing Pediatric Liver Maps

While our healthy pediatric map represents the first map combining cells from multiple healthy donors, there does exist a map from one non-diseased pediatric liver[9] and there is a pediatric diseased liver map from multiple donors with either Biliary Atresia or Choledochal Cysts[61]. We integrated these existing maps with ours to explore study and disease specific differences. The study which includes a healthy pediatric liver[9] enriched for liver immune cells, so while this a valuable resource this map has limited use to understand liver hepatocytes, cholangiocytes, mesenchyme and endothelial cells (Fig S10).

The previously published work on Biliary Atresia focused on differences in cholangiocytes using liver samples from patients with Choledochal Cysts as the comparator[61]. When comparing our healthy pediatric cholangiocytes to cholangiocytes from Biliary Atresia, we found some of the same differences as previously reported (e.g. *IL32* more highly expressed in Biliary Atresia) (Fig. S11B). However, we saw that one key finding, the higher expression of *TNFRSF12A* in Biliary Atresia, may have been driven by a down-regulation in Choledochal Cysts rather than an overexpression in Biliary Atresia (Fig. S11C). When compared to cholangiocytes from healthy pediatric livers, we saw significantly lower expression of *TNFRSF12A* in cholangiocytes from patients with Choledochal Cysts but no difference between Biliary Atresia and healthy pediatric cholangiocytes (Fig. S11C).

Integrating the Biliary Atresia map allowed us to explore the disease specificity of our IFALD results. Comparing healthy pediatric KC like cells to KC like cells from each of the diseases we saw *LY96* specifically overexpressed in IFALD cells and not overexpressed in neither of the other diseases (Fig. S12). In addition, we were able to see a shared signal of MHCI and MHCII overexpression across KC like cells in all three diseases (Fig. S12B).

#### 1 Supplementary Figures

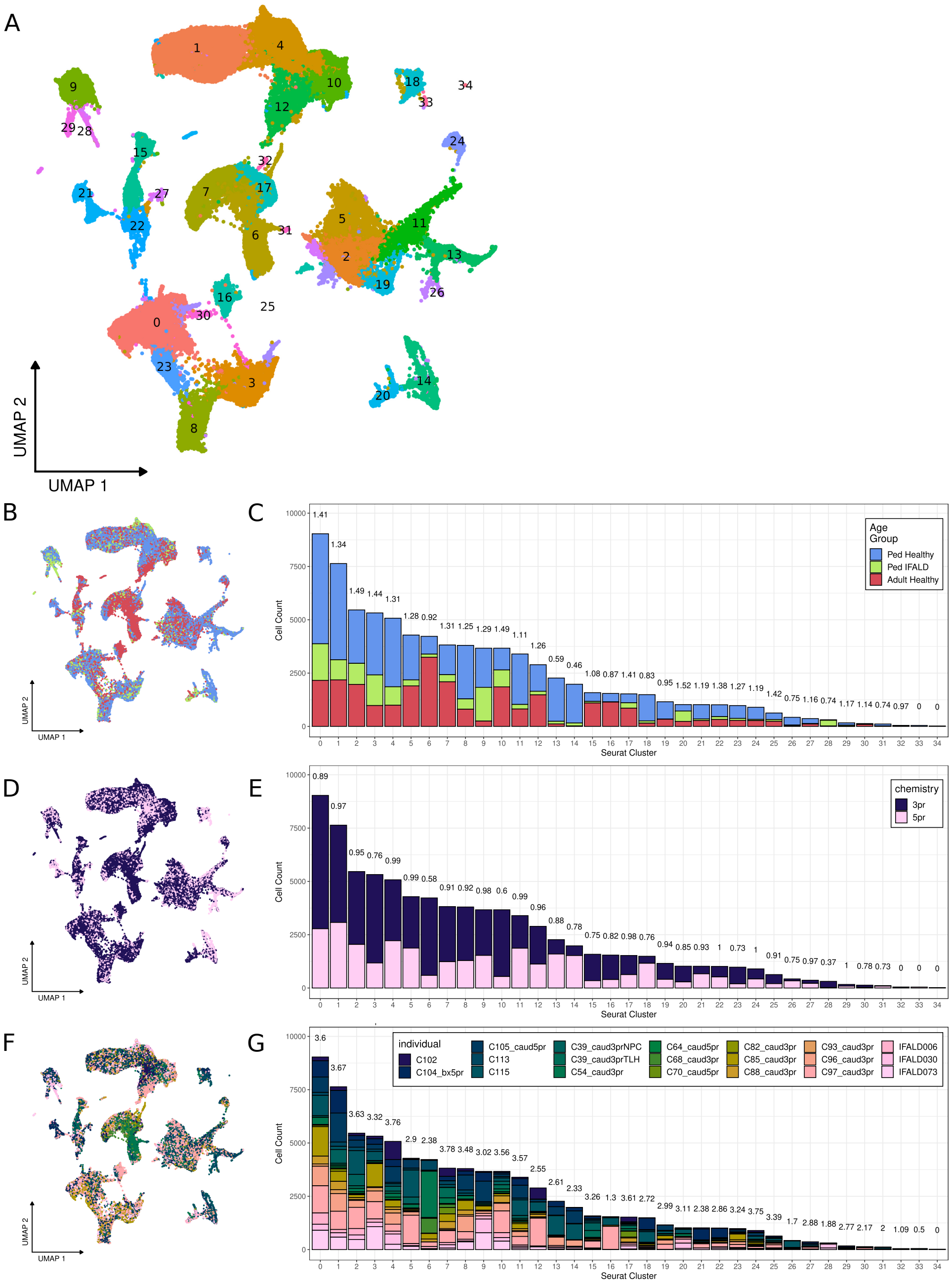


**Figure S1: Datasets are well integrated with high entropy across age, sample type and chemistry in clusters.** A) the liver map UMAP points are colored by clusters. Followed by the UMAP colored by B) age and disease D) Chemistry F) Individual. C-E) Show the contribution of the groups to each cluster found in the map. Number on top of bars is the entropy of the cluster.

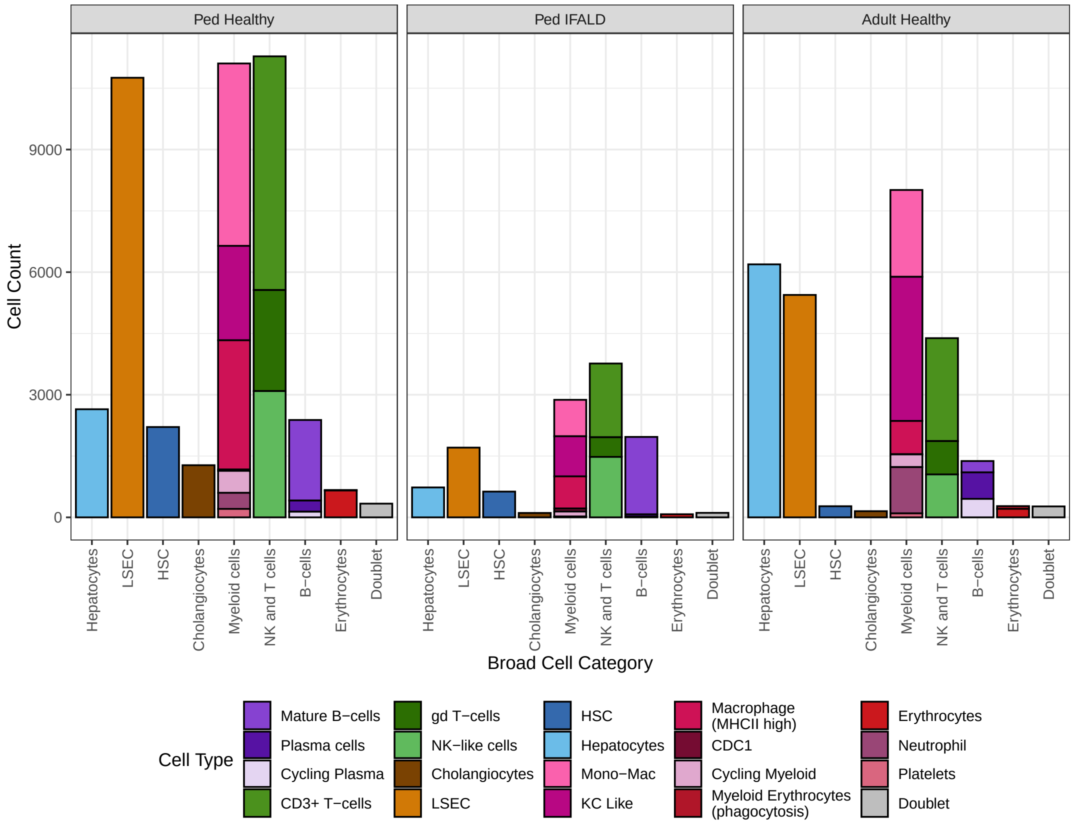


**Figure S2: Cell populations are in similar proportions across age and in IFALD.** Cell counts are grouped broadly by cell category, then split into colored bars of sub populations of cell types.

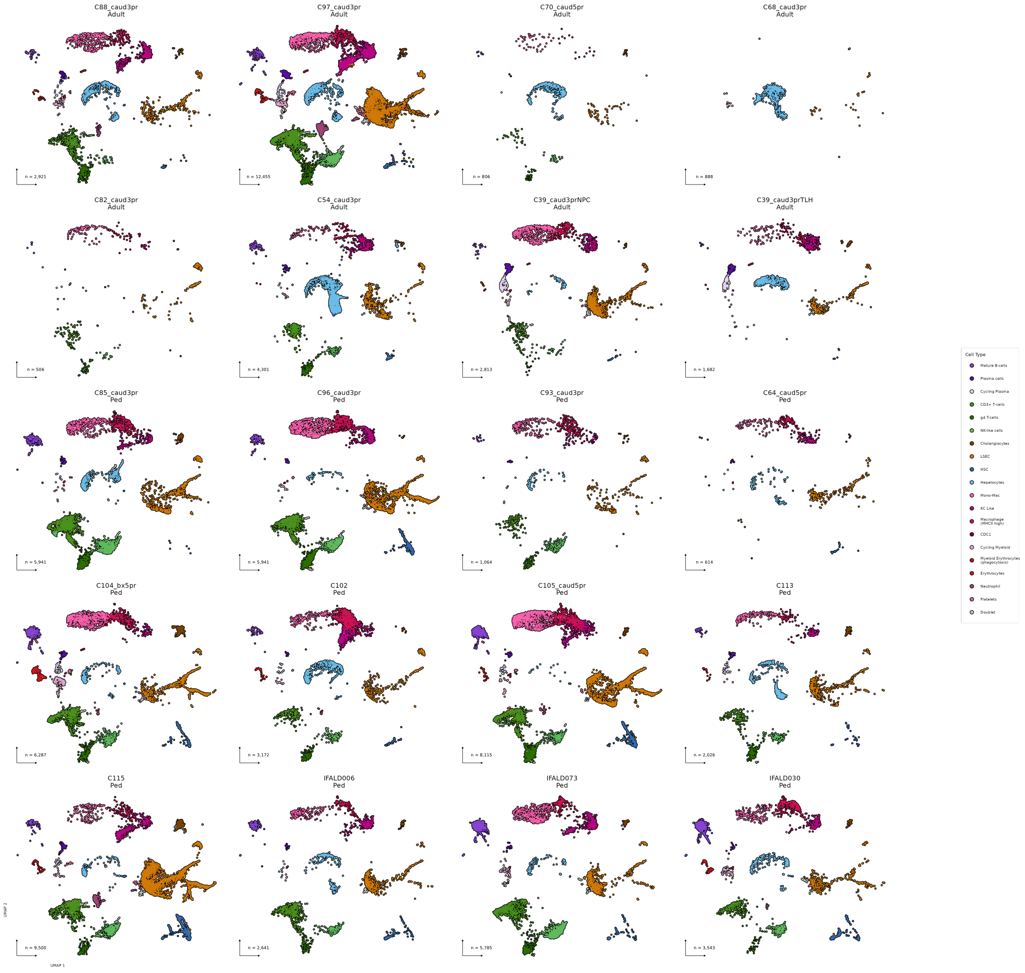


**Figure S3: Cell types are generally present in all individuals.** UMAP of pediatric map split by individual and colored by cell type.

A

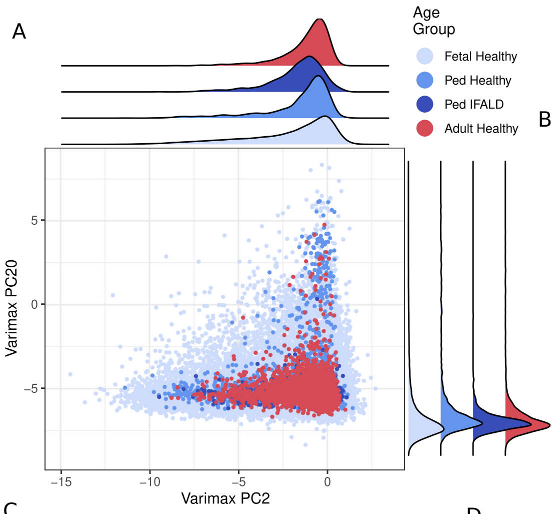


B

**PC2 Top Genes**

| Gene | Score |
|---|---|
| EGR2 | –0.2589401 |
| CCL3L3 | –0.2421634 |
| CXCL3 | –0.2150621 |
| TNF | –0.2097777 |
| IL1B | –0.2081766 |
| CCL4L2 | –0.2067525 |
| IL6 | –0.1982582 |
| CXCL2 | –0.1824378 |
| CCL3 | –0.1783550 |
| CXCL8 | –0.1727274 |

**PC19 Top Genes**

| Gene | Score |
|---|---|
| TMCC2 | 0.3938648 |
| RUNDC3A | 0.2517841 |
| SLC25A37 | 0.1851165 |
| HBA1 | 0.1701209 |
| ARL4A | 0.1680720 |
| TRIM58 | 0.1673931 |
| HBA2 | 0.1621079 |
| HBG2 | 0.1588926 |
| PIM1 | 0.1347608 |
| SLC2A1 | 0.1345923 |

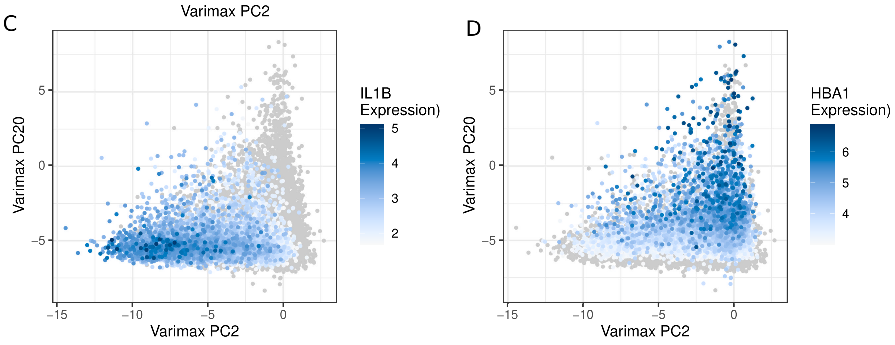


E

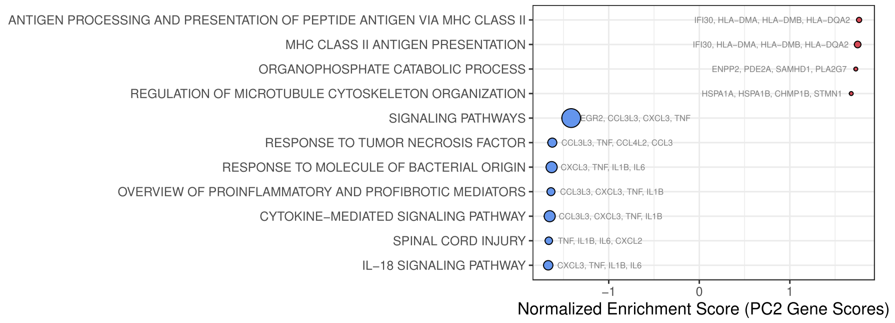


**Figure S4: Varimax PCA captures a factor associated with immune activation.** A) Rotated PC scores for PC2 and PC20. Density plots show the distribution of the scores split by sample age and condition groups. B) The top 10 genes for PC2 and PC20. C-D) Same plot as in A but colored by the expression value of IL1B or HBA1. E) GSEA of the gene associated with PC2.

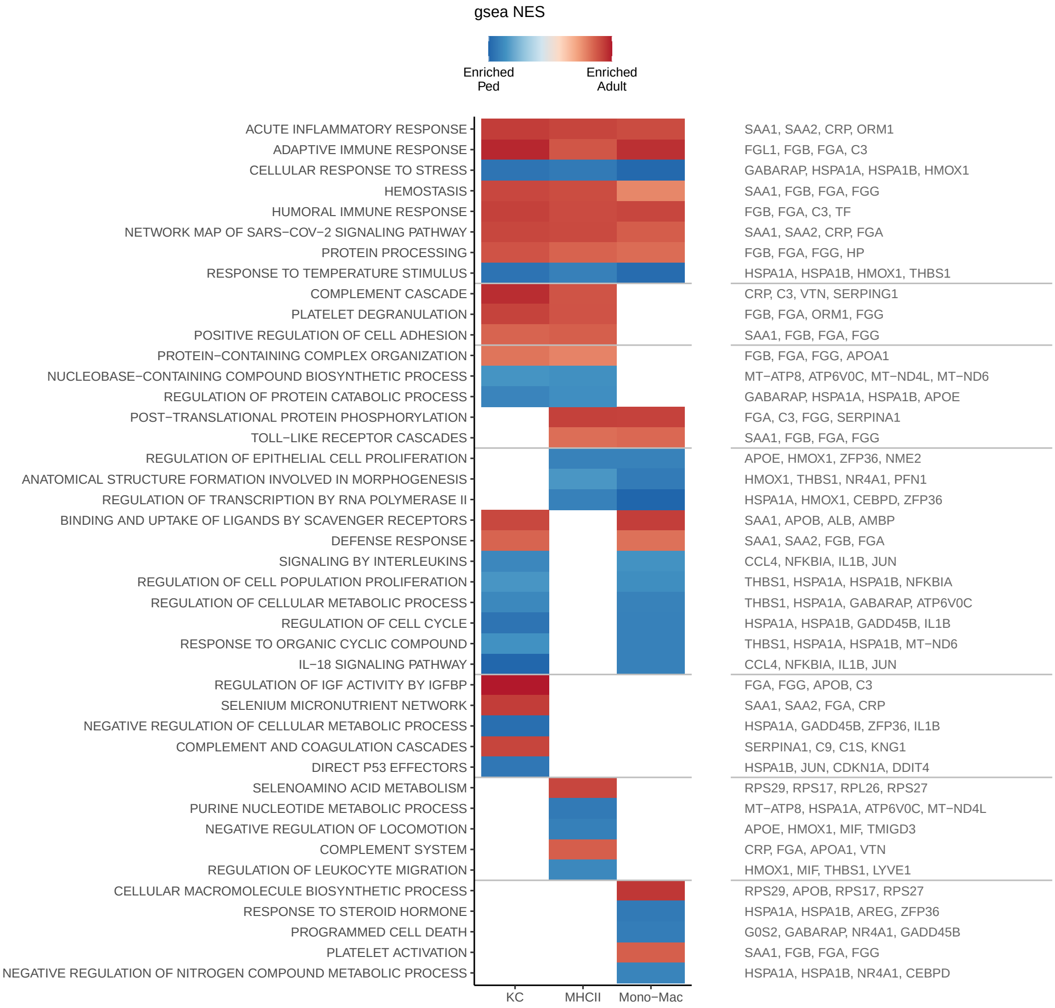


**Figure S5: Pediatric liver myeloid cells express more inflammatory cytokines than adult.** GSEA of genes differentially expressed between pediatric and adult myeloid cell. color of the heat map individual the direction of enrichment of gene sets. Columns are split by myeloid type. Horizontal lines divide genesets into the following categories: enriched in all three myeloid types, enriched in two myeloid types, or enriched in only one myeloid type. Gene names on each row are four random genes from the geneset.

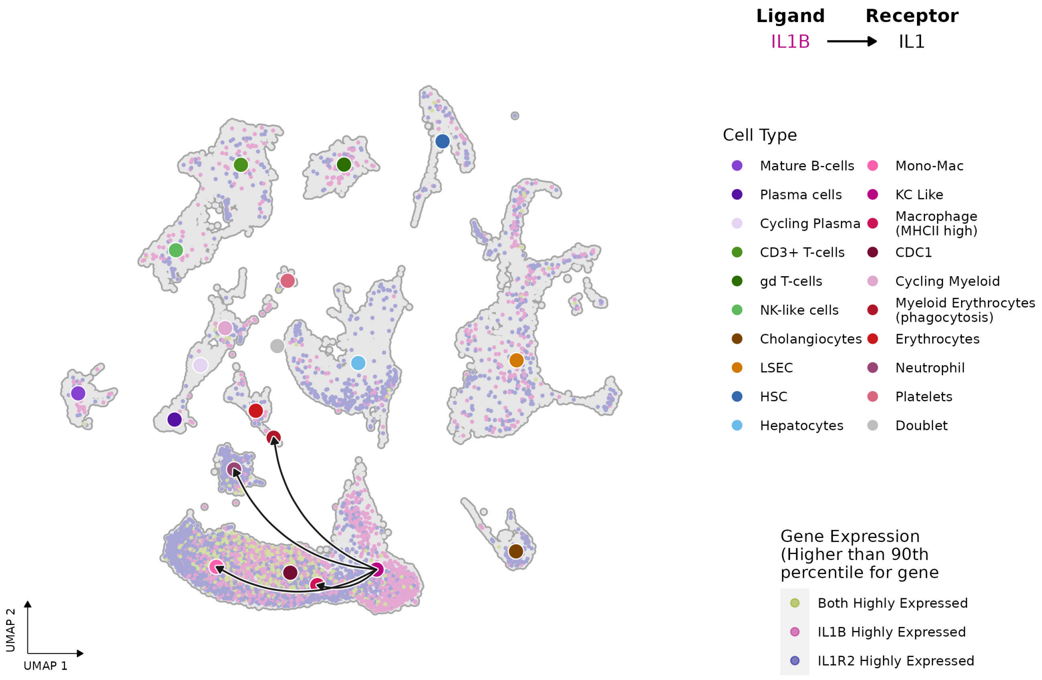


**Figure S6: Predicted cell-cell interactions between KC-like cells and all other cell types through IL1B-IL1 receptor inhibitor.** Individual cells in the UMAP are colored by the expression of the ligand and/or receptor. Arrows on the connecting curves point from ligand to receptor and curves are only shown for significant cell-cell interactions. Larger points which curves connect are colored to indicate the cell population.

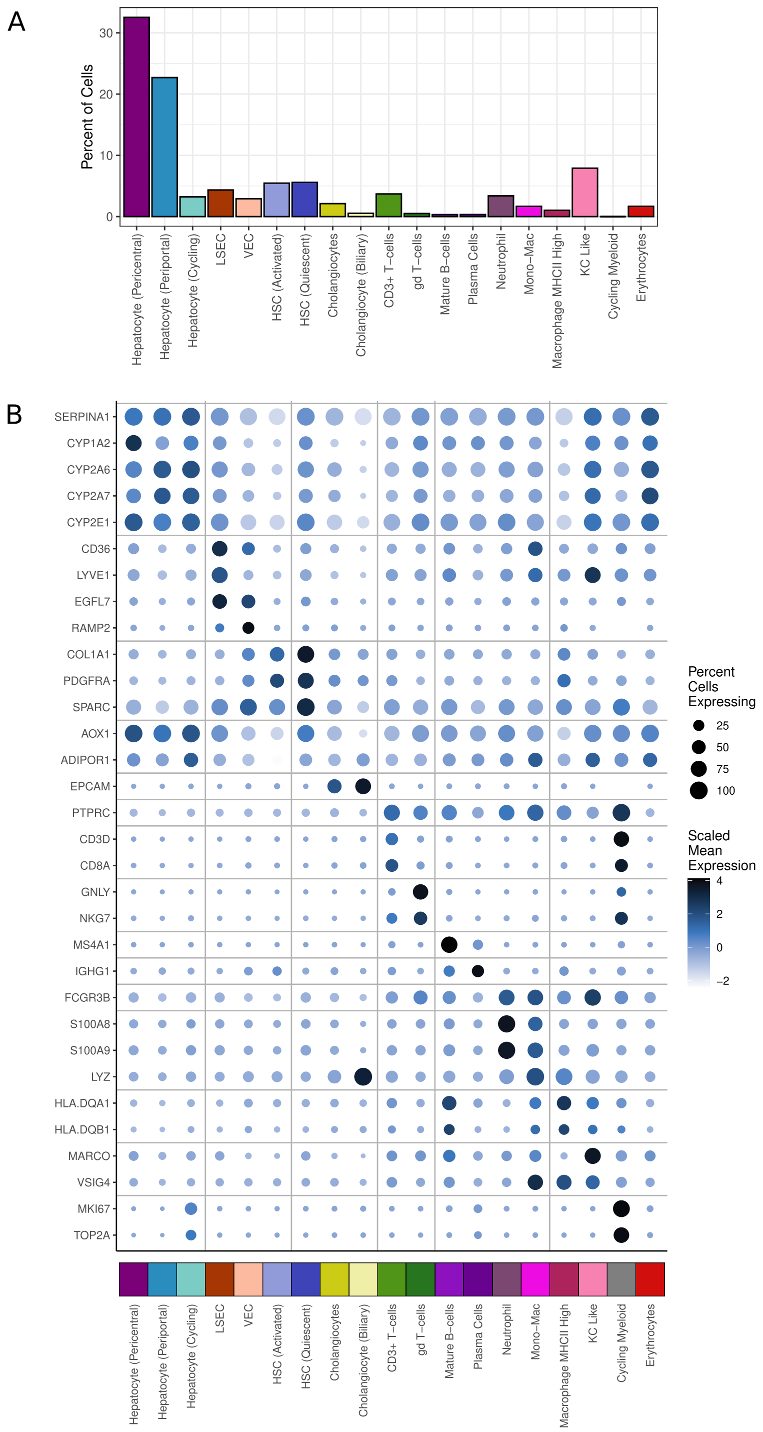


**Figure S7: Xenium spatial transcriptomics of the adult and pediatric liver captures major cell types.** A) Percent of all cells captured with the Xenium of each cell type. B) UMAP of Xenium data colored by assigned cell type. C) Gene expression of key markers used for cell type annotation. The color of the points represents the gene expression level and size represents percent of cells of a type expressing the gene at all.

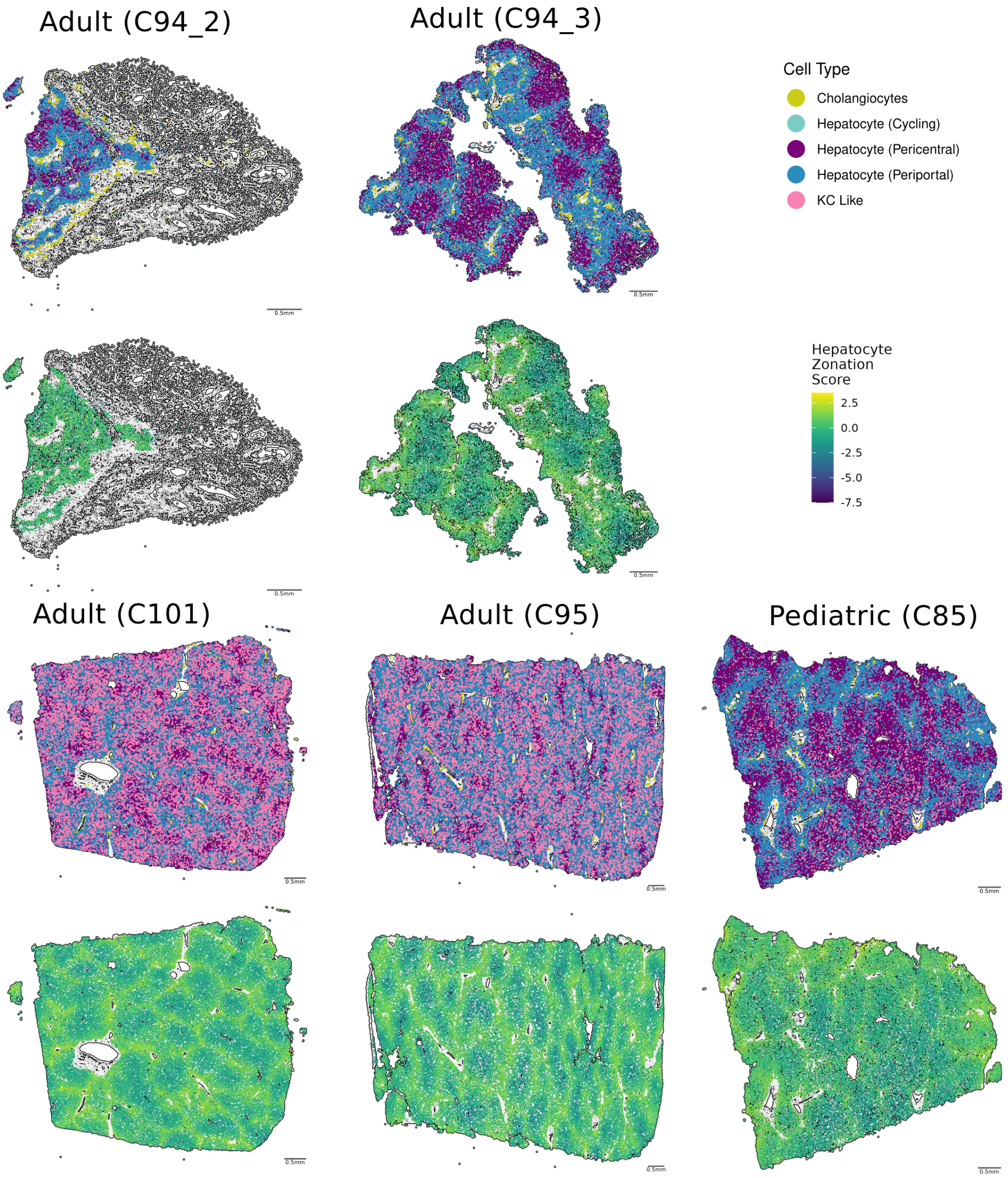


**Figure S8: Xenium spatial transcriptomics of the adult and pediatric liver captures hepatocyte zonation.** Xenium data for each samples, with points representing cells colored by cell type on top and below only hepatocytes are shown with cells colored by zonation score.

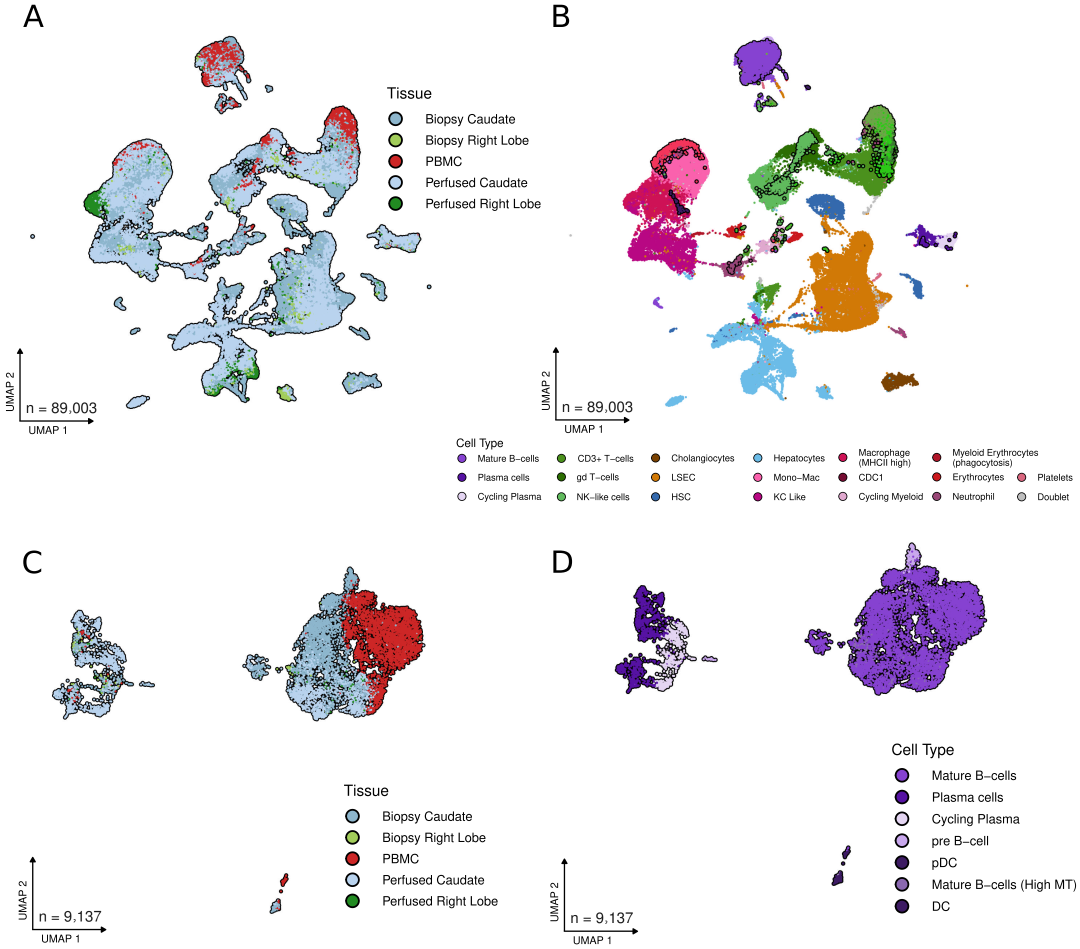


**Figure S9: Integration of PBMC with pediatric liver map.** A) UMAP of integrated PBMC and liver samples with cells colored by sample type. B) Same UMAP colored by cell type with cells from PBMC outlined in black C) UMAP of B cell populations from the PBMC and liver integrated map. Cells are colored by sample type. D) UMAP of B cell populations from the PBMC and liver integrated map. Cells are colored by cell type.

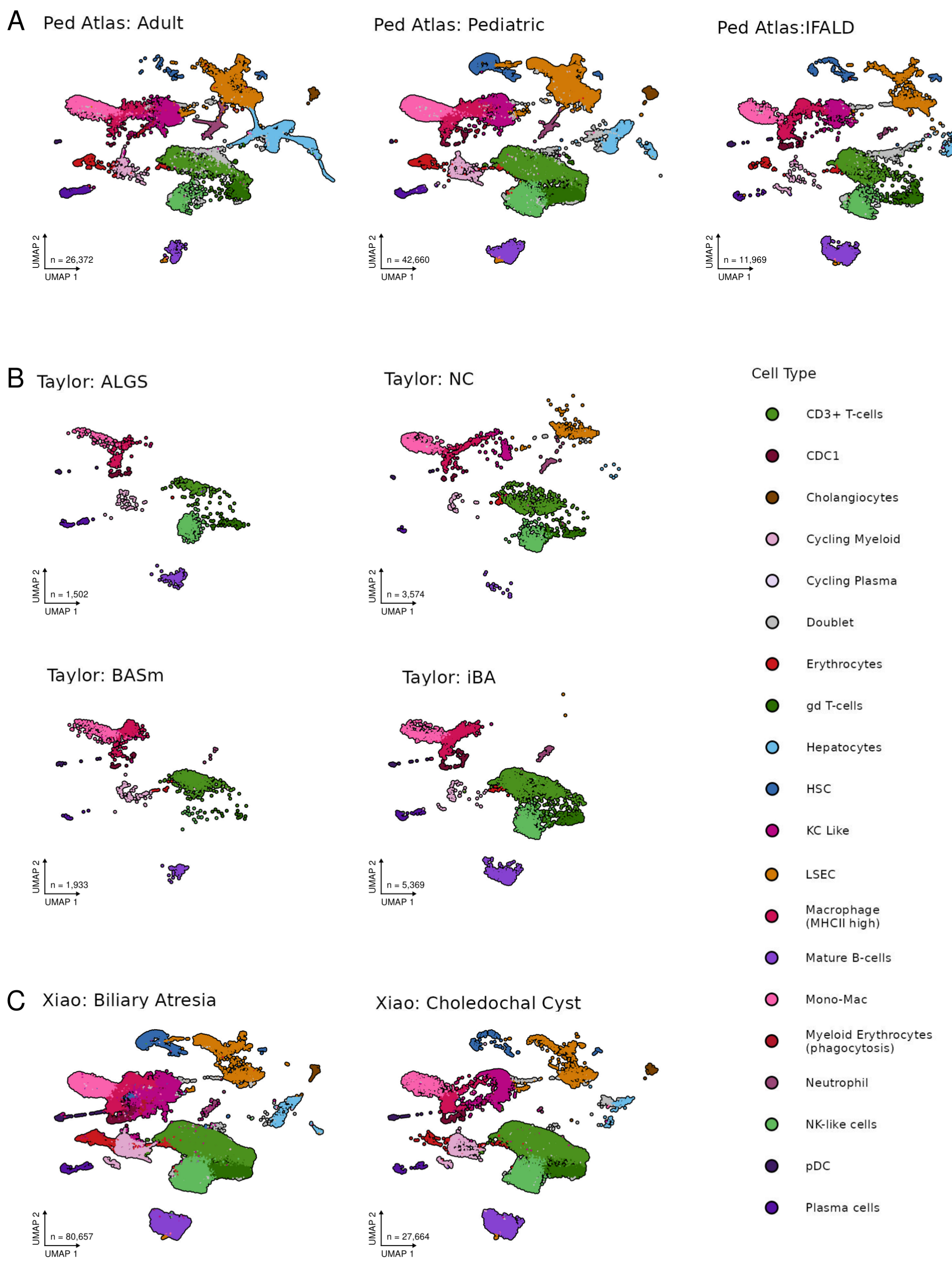


**Figure S10: Cell types are generally present in all conditions with the exception of non-immune cells in the immune enriched Taylor dataset.** UMAP of the pediatric map integrated with previously published pediatric liver data. Map is split by disease and age (i.e. adult liver panel) and coloured by cell type.

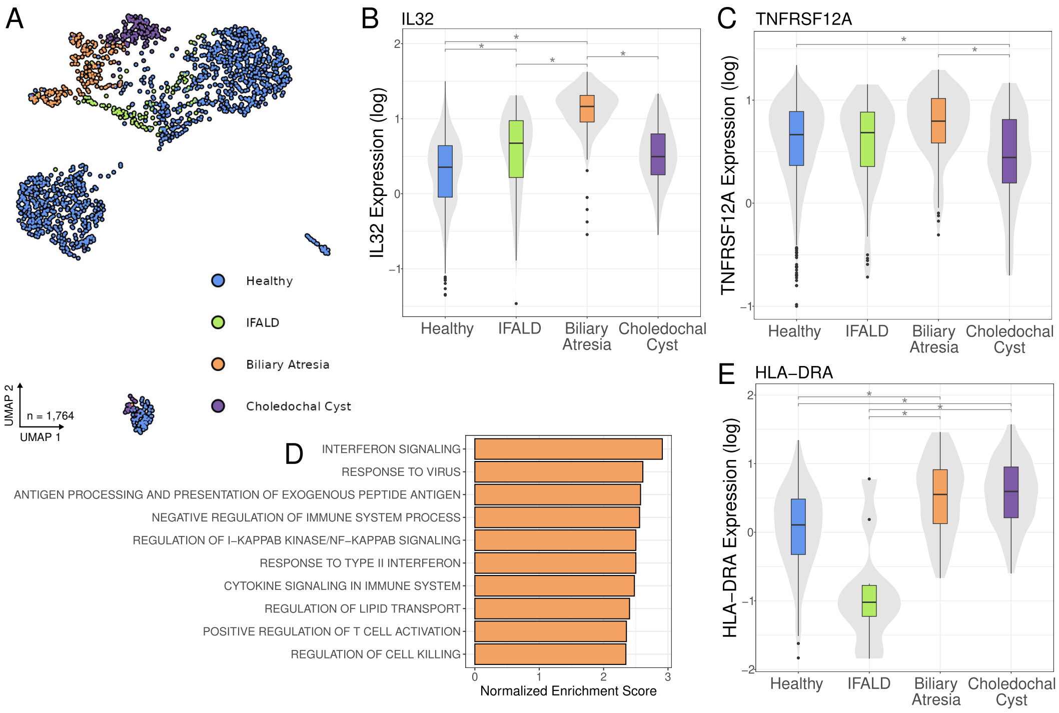


**Figure S11: Pediatric liver cholangiocytes express genes differentially across diseases.** A) Integrated pediatric cholangiocyte UMAP with points coloured by condition. B,C and E) Expression of genes across conditions D) GSEA of differentially expressed genes between healthy pediatric cholangiocytes and cholangiocytes from patients with Biliary Atresia.

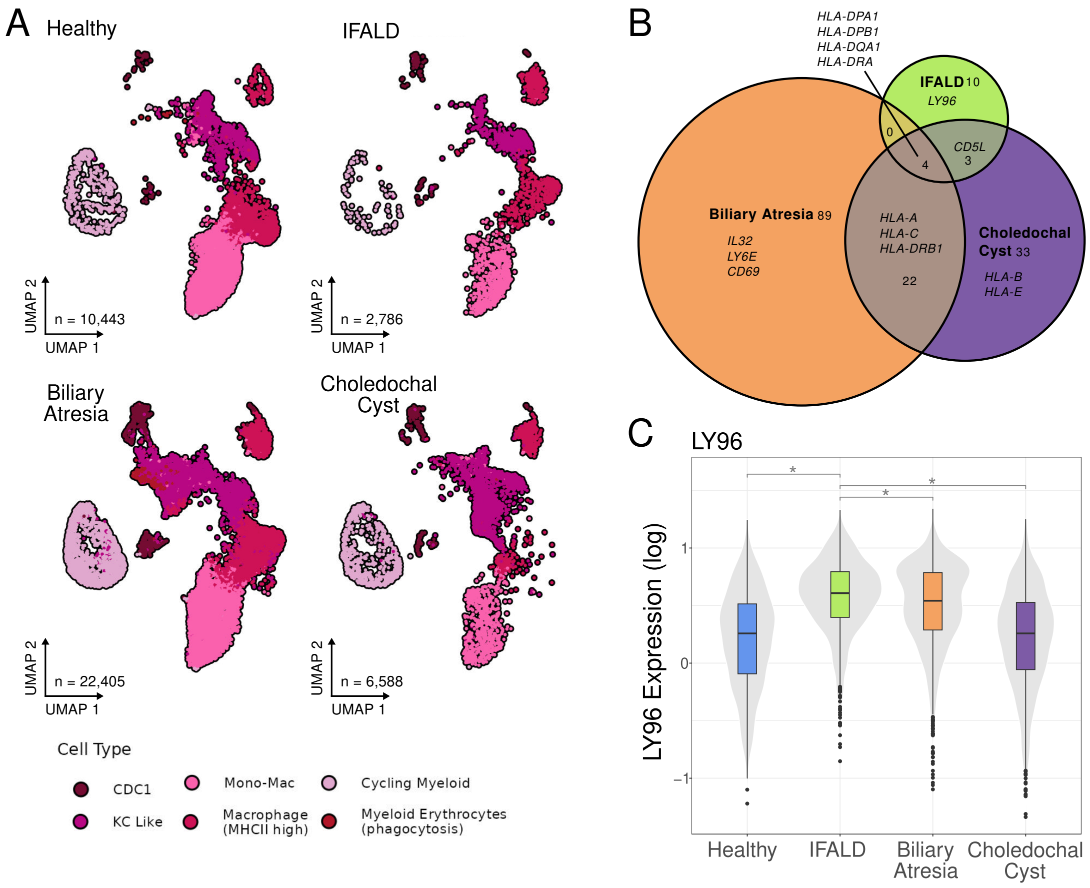


**Figure S12: Pediatric myeloid cells show disease shared and disease specific differential gene expression.** A) UMAPs of pediatric liver myeloid cells split by condition. B) Venn diagram showing the overlap of genes differentially expressed in KC like cells between each disease and to healthy pediatric cells. Only genes upregulated in disease are shown. Numbers indicated the number of genes in each section. C) Expression of LY96 in KC like cells from each condition.

### 2 Supplementary Tables

**Table S1:** scRNA-seq sample information. *NPC: Non-parenchymal cells; TLH: Total liver homogenate; PBMC: Peripheral Blood Mononuclear Cells

| Individual | Dissociation | Sample Type* | Chem | Raw Cell Count | QC Cell Count | Percent Cells Passing QC | Mean gene/ cell | Previously Published (citation) |
|---|---|---|---|---|---|---|---|---|
| C104 | Manual | TLH | 5pr | 19340 | 6287 | 32.51 | 1989 | - |
| C105 | Perfusion | TLH | 5pr | 12332 | 8115 | 65.8 | 2335 | - |
| C85 | Perfusion | TLH | 3pr | 10620 | 5941 | 55.94 | 2006 | - |
| C115 | Manual | TLH | 5pr | 15993 | 9500 | 59.4 | 1911 | - |
| C93 | Perfusion | TLH | 3pr | 1684 | 1064 | 63.18 | 1666 | - |
| C102 | Perfusion | TLH | 5pr | 11071 | 3172 | 28.65 | 2038 | - |
| C113 | Manual | TLH | 5pr | 10285 | 2026 | 19.7 | 1207 | - |
| C64 | Perfusion | TLH | 5pr | 56622 | 614 | 1.08 | 861 | 8 |
| C96 | Perfusion | TLH | 3pr | 7936 | 5941 | 74.86 | 1738 | - |
| IFALD006 | Manual | TLH | 3pr | 8404 | 2641 | 31.43 | 1172 | - |
| IFALD030 | Manual | TLH | 3pr | 6550 | 3543 | 54.09 | 2276 | - |
| IFALD073 | Manual | TLH | 3pr | 10217 | 5785 | 56.62 | 2109 | - |
| IFALD073 | PBMC | PBMC | 3pr | 8689 | 8002 | 92.09 | 1969 | - |
| C82 | Perfusion | TLH | 3pr | 12821 | 506 | 3.95 | 1588 | - |
| C70 | Perfusion | TLH | 5pr | 10000 | 806 | 8.06 | 2269 | 8 |
| C97 | Perfusion | TLH | 3pr | 21057 | 12455 | 59.15 | 1802 | - |
| C68 | Perfusion | TLH | 3pr | 65540 | 888 | 1.35 | 544 | 8 |
| C39 | Perfusion | NPC | 3pr | 7604 | 2813 | 36.99 | 923 | - |
| C39 | Perfusion | TLH | 3pr | 6515 | 1682 | 25.82 | 788 | 1,8 |
| C54 | Perfusion | TLH | 3pr | 19349 | 4301 | 22.23 | 861 | 8 |
| C88 | Perfusion | TLH | 3pr | 11111 | 2921 | 26.29 | 1508 | - |

**Table S3:** Spatial transcriptomic sample information

| Individual | Tissue | Structures of note | Sex | Age (years) | Median Transcripts | Median Genes | Cells |
|---|---|---|---|---|---|---|---|
| C105 | Right Lobe | parenchyma | F | 7-12 | 399 | 66 | 66295 |
| C85 | Caudate | parenchyma | M | 7-12 | 258 | 61 | 65291 |
| C94_2 | Caudate | bile duct hamartoma | F | 38 | 91 | 44 | 37334 |
| C94_3 | Caudate | parenchyma | F | 38 | 572 | 77 | 36179 |
| C94_4 | Caudate | parenchyma | F | 38 | 554 | 76 | 47873 |
| C95 | Caudate | parenchyma | F | 47 | 361 | 63 | 123326 |
| C101 | Caudate | parenchyma | M | 43 | 328 | 59 | 108999 |

**Table S5:** List of anitbodies

| Target | Source | Identifier |
|---|---|---|
| CD3 | Thermofisher | 414-0032-80 |
| CD45 | BioLegend | 304824 |
| CD68 | BioLegend | 333821 |
| CD14 | BioLegend | 325603 |
| IL1-β | Thermofisher | 51-7018-42 |
| CD68 | Agilent | PG-M1 |
| MD-2 | Abcam | ab24182 |
